# The Quorum Sensing regulated sRNA Lrs1 is involved in the adaptation to low iron in *Pseudomonas aeruginosa*

**DOI:** 10.1101/2024.12.09.627364

**Authors:** Dimitra Panagiotopoulou, Natalia Romo Catalán, Max Wilcox, Nigel Halliday, Paolo Pantalone, James Lazenby, Miguel Cámara, Stephan Heeb

**Author notes:** **Corresponding Authors:** Dimitra Panagiotopoulou, Stephan Heeb.

## Abstract

Iron is an essential nutrient for microbial growth. The opportunistic pathogen *Pseudomonas aeruginosa* can survive under diverse conditions, including iron-depleted environments with the aid of small non-coding RNAs (sRNAs). *P. aeruginosa* also uses three quorum sensing (QS) systems Las, Rhl, and Pqs to coordinate virulence and infection establishment at the population level. This study links the sRNA Lrs1, which is located within the promoter of the Pqs biosynthetic operon *pqsABCDE,* to iron uptake regulation in the *P. aeruginosa* strain PAO1-L. Transcriptomics and phenotypic assays indicate that Lrs1 downregulates the production of the siderophore pyochelin but not pyoverdine, and that *lrs1* regulation itself is dependent on iron availability. Although Lrs1 has been implicated in a positive feedback loop with the transcriptional regulator LasR in the strain PA14, the present findings indicate that this is not the case in PAO1-L in the tested conditions. Transcription of Lrs1 is dependent on quorum sensing, predominantly on RhlR with an auxiliary effect by PqsE. Furthermore, the Pqs system and phenazine production are modulated by Lrs1 only under iron limitation. This study identifies Lrs1 as a new QS-dependent post-transcriptional regulator of iron uptake and virulence highlighting its importance in environmental adaptation in *P. aeruginosa*.

**Graphical abstract:** 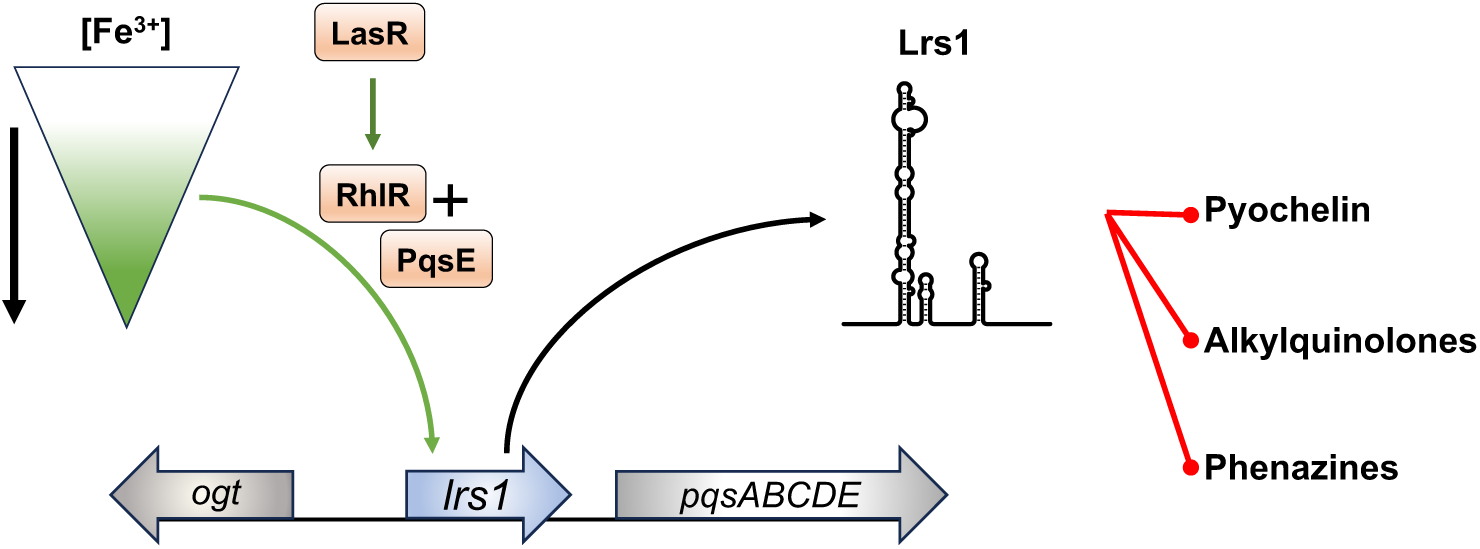

## Introduction

*Pseudomonas aeruginosa* is an opportunistic pathogen with the ability to colonise diverse environments. It is one of the most common healthcare associated pathogens, causing antibiotic resistant infections and a major morbidity risk factor for cystic fibrosis (CF) patients (CDC, 2019; Malhotra, Hayes and Wozniak, 2019). The pathogenesis of *P. aeruginosa* is controlled at the population level via the coordinated action of three quorum sensing (QS) systems, the Las, Rhl, and Pqs systems. The Las and the Rhl QS systems, rely on the production and sensing of *N*-acyl-homoserine lactones as their cognate signals. The complex LasR - *N*-(3-oxododecanoyl)-L-homoserine lactone (3OC_12_-HSL) activates the expression of genes involved in biofilm formation and virulence, such as that encoding the secreted elastase LasB (Williams and Cámara, 2009). Homeostasis in the activation of the Las system is preserved by the negative transcriptional regulator RsaL which binds to the bidirectional promoter of *rsaL*-*lasI* repressing both its own expression and that of the 3OC_12_-HSL synthetase gene *lasI* (Rampioni *et al*., 2007). The RhlR transcriptional regulator forms a complex with the *N*-butanoyl-L-homoserine lactone (C_4_-HSL) controlling the production of virulence factors including hydrogen cyanide, pyocyanin, and determinants of biofilm formation (Williams and Cámara, 2009). The third QS system, known as the Pqs system, is driven by 2-alkyl-4(1*H*)-quinolones (AQ) signal molecules. For the biosynthesis of these molecules, chorismate is converted to anthranilate via the action of PhnAB or TrpEG and subsequently, to AQs via the products of the *pqsABCDE* operon and *pqsH* (Farrow and Pesci, 2007). The PqsR transcriptional regulator binds either 2-heptyl-4-hydroxyquinoline (HHQ) or 2-heptyl-3-hydroxy-4-quinolone also referred to as the *Pseudomonas* quinolone signal (PQS), and enhances the expression of the *pqs* operon, as well as the production of multiple virulence traits including pyocyanin and the release of eDNA contributing to biofilm architecture (Xiao *et al*., 2006; Rampioni *et al*., 2010). However, RhlR negatively regulates both the *pqsABCDE* operon and the *pqsR* gene (Xiao, He and Rahme, 2006). Besides its role in QS, the PQS molecule has iron chelating properties, restricting access to environmental iron by competing bacteria, further enhancing the adaptation of *P. aeruginosa* infections in a polymicrobial environment (Bredenbruch *et al*., 2006; Diggle *et al*., 2007).

The QS systems can also respond to environmental signals at the post-transcriptional level through the action of small non-coding RNAs (sRNAs). These sRNAs can act by base pairing with target mRNAs altering their translational rate or stability, and these interactions may be dependent on the RNA chaperone Hfq (Wagner and Romby, 2015). Under iron limitation, the expression of the Pqs system is increased by the indirect action of the Prrf1 and Prrf2 sRNAs (Oglesby *et al*., 2008). These two redundant sRNAs repress *antR*, encoding the regulator of anthranilate degradation, redirecting anthranilate, the alkyl-quinolone precursor, towards the biosynthesis of PQS. This post-transcriptional regulation is antagonised by the Carbon Catabolism Regulator sRNA CrcZ, allowing for more anthranilate to de degraded by products of the AntR-induced genes *antABC* and *catBCA* towards the TCA cycle (Sonnleitner, Prindl and Bläsi, 2017). The sRNA PhrS positively affects the Pqs QS system from two fronts; it increases the translation rate of *pqsR* while repressing that of *antR,* hence more anthranilate is directed to PQS production (Sonnleitner *et al*., 2011; Gebhardt *et al*., 2023).

In infection sites such as the CF lung, pathogens compete for access to iron (Hunter *et al*., 2013; Konings *et al*., 2013). *P. aeruginosa* possesses multiple systems for iron uptake from the environment aiding to the establishment of acute and chronic infections (Cornelis and Dingemans, 2013; Konings *et al*., 2013). *P. aeruginosa* produces two siderophores, pyoverdine and pyochelin with high and low affinity for iron, respectively (Brandel *et al*., 2012; Cornelis and Dingemans, 2013). It is hypothesised that the low iron affinity pyochelin is produced first, while the more energy-demanding pyoverdine is reserved for iron scarcity (Dumas, Ross-Gillespie and Kümmerli, 2013; Cunrath *et al*., 2020). Pyochelin biosynthesis, like PQS, starts with chorismic acid. Chorismate is converted to salicylate followed by condensation with two cysteines (Reimmann *et al*., 1998). The pyochelin biosynthetic operons *pchDCBA* and *pchEFGHI*, along with the cognate Fe(III)-pyochelin receptor gene *fptA* are induced by the transcriptional regulator PchR bound to the ferripyochelin complex (Michel *et al*., 2005). When iron is low, the sRNAs PrrF1,2 negatively regulate the mRNAs of non-essential iron-containing proteins, preserving iron for essential processes (Wilderman *et al*., 2004). When iron is present, both transcription of Prrf1,2 and production of the siderophores are repressed by the iron regulator Fur, while the iron storage protein BfrB is upregulated thus maintaining iron homeostasis (Wilderman *et al*., 2004; Cornelis, Matthijs and Van Oeffelen, 2009).

The sRNA Lrs1 transcribed within the *pqsA* promoter region has been shown to be involved in a positive regulatory feedback loop with LasR, in surface-attached cells of the *P. aeruginosa* strain PA14 (Chuang *et al*., 2019). Deletion of *lrs1* resulted in reduced transcript levels of *lasR* in both planktonic and surface-attached cells, with a more pronounced effect in the latter. This deletion also abolished elastase production in planktonic cultures. However, in contrast to these observations, a previous RNA-seq analysis of the same Δ*lrs1* deletion mutant in planktonic cells revealed that neither the expression of *lasR* nor *lasB* were altered compared to the wild type (Wurtzel *et al*., 2012). In this study we show how we have independently identified *lrs1* in the *P. aeruginosa* PAO1-L strain through an RNA-seq analysis and how we sought to address the discrepancies mentioned above by investigating the role that Lrs1 has in the QS systems of this this strain. We found that in PAO1-L, Lrs1 does not significantly impact on the Las and Rhl QS systems in the conditions tested. However, the Pqs QS system seems to be conditionally affected, with no observed effect when grown in LB but a downregulation by Lrs1 in iron scarcity. Furthermore, we found that Lrs1 is involved in the regulation of iron acquisition systems, including the two siderophores pyochelin and pyoverdine, and that this regulation is dependent on iron availability. This regulation may be due to the participation of Lrs1 in balancing metabolic flow, deprioritising pyochelin, PQS, and phenazine production in favour of central metabolism and aromatic amino acid biosynthesis. This is the first demonstration of the differential regulation of these siderophores by a post-transcriptional regulator and a regulatory connection between the Pqs QS system and pyochelin, offering novel mechanistic insight into the regulation of iron acquisition in *P. aeruginosa*.

## Results

### *Lrs1* appears in multiple transcript forms

Lrs1 is an sRNA that lies upstream of the *pqsA* coding sequence. BlastN analysis of Lrs1 showed its existence only within *P. aeruginosa*, with the shortest size of a homologue being 190 nt long in strain PA14 (Wurtzel *et al*., 2012). In our lab, Lrs1 was independently identified in an RNA-seq analysis carried out in strain PAO1-L after 8 h and 12 h of incubation in LB (Supplementary Figure 1 A). In the *pqsA* promoter (P*_pqsA_*) region, two transcription start sites (TSS) have been identified, TSS1 and TSS2 (Dötsch *et al*., 2012). Transcription from TSS2 appeared to continue beyond the 190 nt end point of *lrs1* suggesting that Lrs1 may exist as a longer transcript (Supplementary Figure 1 B). To investigate this possibility, Northern blot was conducted in the PAO1-L wild type (WT) strain after 12 h of incubation in LB to match the time point of the RNA-seq, and compared to the bands produced by an *lrs1* deletion mutant (Figure 1 B). Indeed, more than one band was observed in the PAO1-L compared to the *lrs1* mutant, suggesting the presence of transcripts of different sizes driven by the TSS2 promoter. The bands observed in Δlrs1 did not present the same pattern as in the WT. Since *lrs1* was only partially deleted and the Northern blot probe binds to a further 52 nt, these bands are products transcribed from TSS2 as deletion of the entire *lrs1* completely eliminated these bands (Supplementary Figure 1 D). The presence of these bands indicate that transcription from TSS2 had not been affected by the deletion of *lrs1.* Detection of Lrs1 in PA14 also revealed the same bands as in the PAO1-L WT, suggesting that Lrs1 is present in multiple forms in *P. aeruginosa*, and that this is not strain-specific (Figure 1 C).

**Figure 1.**
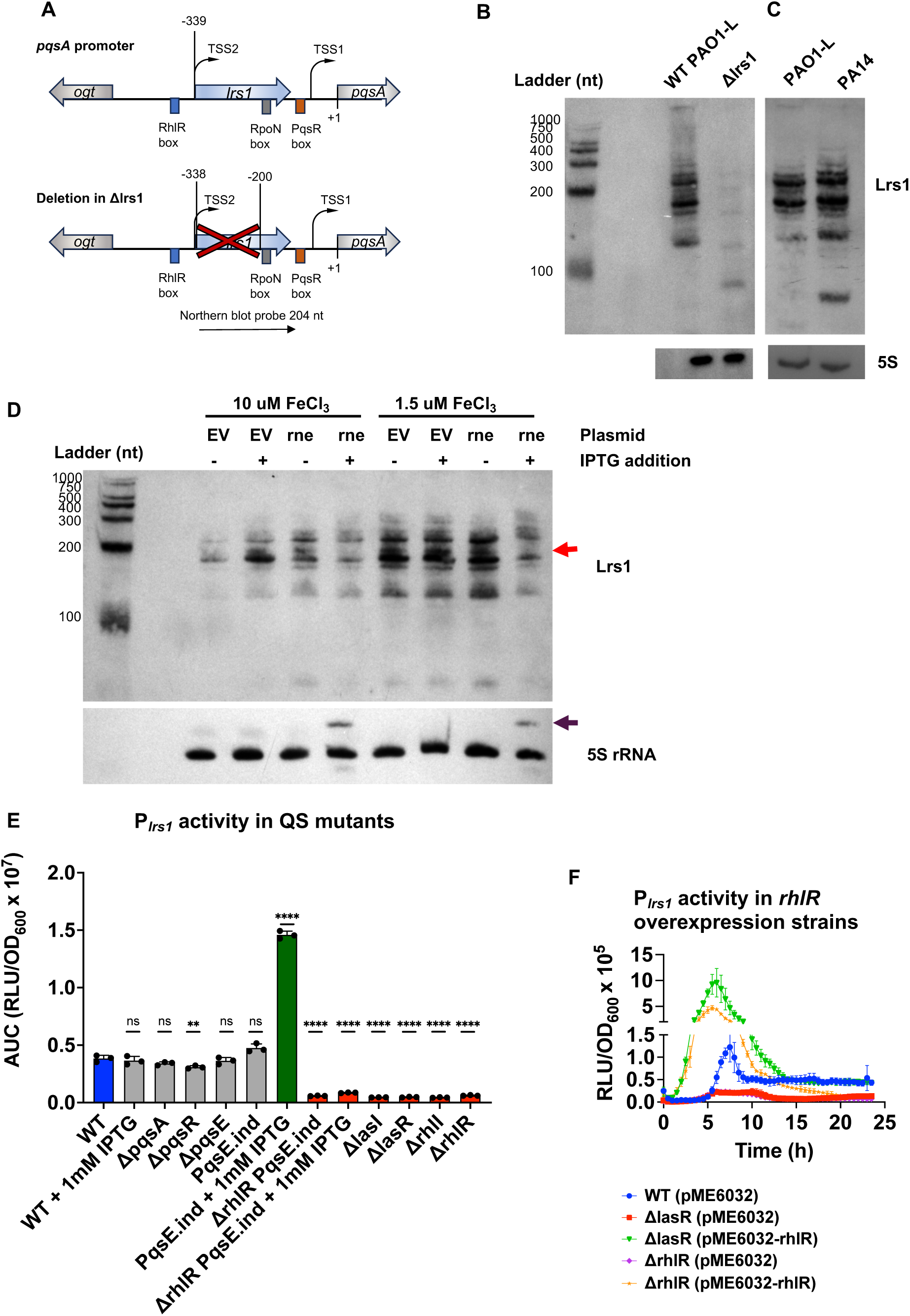
Lrs1 abundance is dependent on RNase E and the *lrs1* promoter activity of the three QS systems. A) Construction of *lrs1* deletion mutant. Schematic representation of the intergenic region between *pqsA* and the upstream *ogt* gene showing the position of *lrs1* starting at TSS2, 339 bp upstream of the *pqsA* translational start site (+1). The positions of RhlR, PqsR and RpoN binding boxes are also depicted with rectangles. Of the 190 bp of *lrs1*, only the first 138 bp were deleted, leaving the TSS2 unaffected. This was done to preserve the RpoN binding site found in the *pqsA* promoter (Shao *et al*., 2018). B) Northern blot of Lrs1 in PAO1-L after 12 h of incubation in LB. C) Northern blot of Lrs1 in PAO1-L and PA14 after 12 h of incubation in LB. D) RNase E is involved in processing and abundance of Lrs1. Northern blot of Lrs1 from cells grown at 10 and 1.5 uM iron. Samples were collected 16 h post-inoculation in M9 medium with glucose and the iron concentration indicated. EV: empty vector, rne: pBx-sgRNA-rne expressing plasmid, constitutively transcribing the guide RNA for the RNase E gene *rne*. IPTG addition induces the expression of the catalytically inactive (dead) Cas9 from the P*_lac_* inducible promoter. The *rne* gene is knocked down and the levels of RNase E are reduced in the cells transcribing the sgRNA targeting *rne* from pBx-sgRNA-rne plasmid. Where indicated, IPTG was added at 1 mM final concentration at the beginning of the culture. 5S rRNA was used as loading control. Red arrow indicates the lack of one Lrs1 band in the IPTG-induced expression of dCas9 in the pBx-sgRNA-rne harbouring strain. Purple arrow represents the presence of a higher molecular weight band of 5S rRNA in induced strains with the pBx-sgRNA-rne plasmid, indicative of loss of 5S rRNA processing due to RNase E knockdown. E) Transcriptional activity of the *lrs1* promoter (P*_lrs1_*) in different QS mutants. Approximately 500 bp upstream of *lrs1* were cloned upstream of the bioluminescence reporter operon *luxCDABE* and promoter activity was measured over 24 h in PAO1-L WT and QS mutants in LB. Luminescence was measured in Relative Luminescence Units (RLU) and was normalised by growth (OD_600_). The Area Under the Curve (AUC) of each curve has been plotted here. Error bars represent the standard deviation of three biological replicates with three technical replicates. One-way ANOVA was used as statistical test. ns: not significant. * : p-value ≤ 0.05, ** : p-value ≤ 0.01, *** : p-value ≤ 0.001, **** : p-value ≤ 0.0001. F) Transcriptional activity of P*_lrs1_* in strains overexpressing *rhlR*. Error bars represent three biological replicates with two technical replicates.

### Abundance of Lrs1 is dependent on RNase E

Often, sRNAs are transcribed as longer precursor transcripts before they are processed post-transcriptionally by RNase E to adopt their shorter, mature forms. RNase E preferentially cuts RNAs in single-stranded regions that are enriched with poly-U, and in *Salmonella typhimurium*, RNase E recognises the pattern RNWUU where R is G/A, N any nucleotide, and W is A/U (Chao *et al*., 2017). Looking at the predicted secondary structures of Lrs1 and the long 5’UTR starting from TSS2, three single-stranded regions contain motifs resembling recognition sites for RNase E Supplementary Figure 2). Therefore, it was hypothesised that RNase E may be involved in the processing of Lrs1.

To explore this hypothesis, RNase E was depleted from the cells using a CRISPR interference approach (CRISPRi) previously developed for *P. aeruginosa* (Tan, Reisch and Prather, 2018) and Northern blots were performed to evaluate its effect on Lrs1. The experiment was conducted in both high and low iron availability, as a connection between Lrs1 and iron is shown later in this study.The abundance of Lrs1 was significantly reduced in RNase E-depleted cultures in low iron conditions, and less so in high iron (Figure 1D). Although reduced RNase E activity often results in increased abundance of RNAs in some cases, it may also have the opposite effect (Stead *et al*., 2011; Mackie, 2013). Additionally, at least one of the bands of Lrs1 disappears in the *rne-*repressed cells compared to cells carrying the empty vector, implicating RNase E in the post-transcriptional processing of Lrs1. In conclusion, RNase E is essential for normal levels and post-transcriptional processing of Lrs1.

### *Lrs1* is regulated by the three QS systems of P. aeruginosa *PAO1-L*

Lrs1 overlaps with the *pqsA* promoter and its transcription is initiated from the distal transcriptional start site of *pqsA*, the TSS2 (Figure 1A). However, little is known on the effect of the three QS systems on the regulation of transcription from TSS2, besides the positive regulation by LasR and RhlR (Wurtzel *et al*., 2012; Brouwer *et al*., 2014). To this end, the 500 bp upstream of TSS2, including the predicted promoter itself, named here as the *lrs1* promoter (P*_lrs1_*) was transcriptionally fused to the *luxCDABE* operon. The activity levels of the promoter were monitored throughout growth in different QS mutants and were compared to the WT.

The activity of the *lrs1* promoter was impacted by deletions in the Las and Rhl but not the Pqs QS systems (Figure 1 E, Supplementary Figure 3). Deletion of *lasR, lasI, rhlR,* or *rhlI* silenced the promoter (Supplementary Figure 3 C, D). Considering the positive regulation of *rhlR* by LasR (Latifi *et al*., 1996), it was hypothesised that the effect of *lasR* deletion may be indirect, through diminished levels of RhlR. To investigate this, both *lasR* and *rhlR* mutants were complemented with a vector overexpressing *rhlR*.

Indeed, overexpression of *rhlR* significantly increased the activity of P*_lrs1_*, including in the *lasR* mutant, indicating that RhlR is the primary regulator of the promoter and the LasR effect is indirect (Figure 1 F). It has been suggested that the effector protein of the Pqs QS system, PqsE, and RhlR physically interact, and that interaction alters the DNA binding affinity of RhlR thus affecting its regulatory activity (Borgert *et al*., 2022; Feathers *et al*., 2022). For this reason, the effect of *pqsE* on P*_lrs1_* was also tested. Deletion of *pqsE* did not impact activity of the promoter (Figure 1 E, Supplementary Figure 3 A). In contrast, overexpression of *pqsE* from an IPTG-inducible mutant, significantly increased the activity levels of P*_lrs1_*, an effect lost upon deletion of *rhlR* in this mutant (Figure 1 E, Supplementary Figure B). In conclusion, the *lrs1* promoter is regulated by the Las and primarily by the Rhl QS systems, with PqsE having a positive yet auxiliary impact on this promoter in PAO1-L.

### *Lrs1* does not participate in the regulation of the QS systems

Considering the location of *lrs1*, it is plausible that a regulatory link between Lrs1 and the *pqs* operon may exist. The activity of a *pqsA-luxCDABE* reporter was not affected at the transcriptional but only slightly at the translational level when *lrs1* was overexpressed (Supplementary Figure 4 A, B, C, D). Quantification of PQS, HHQ, and HQNO in culture supernatants overexpressing *lrs1*, showed no significant difference when compared to the strain with the empty vector (Supplementary Figure 4 F, G, H, I). It was noted that deletion of *lrs1* reduced the production of HHQ while increasing the levels of its precursor anthranilate, but this phenotype could not be complemented by the overexpression of *lrs1* from the pKH6 plasmid. Both the WT and Δlrs1 were whole genome sequenced with no single polynucleotide polymorphisms (SNPs) detected between the two strains suggesting that the HHQ reduction was attributed solely on the *lrs1* deletion. Additionally, the deletion of *lrs1* impacted the transcriptional output of P*_pqsA_* (Supplementary Figure 4 J, K), indicating that reduced HHQ production may be due to reduced activity from the *pqsA* promoter and not an effect attributed to Lrs1. Therefore, Lrs1 does not seem to have a major impact in the regulation of the *pqs* operon under the tested conditions.

Previous work on Lrs1 found a positive regulatory link between Lrs1 and the Las QS system in PA14 (Wurtzel *et al*., 2012; Chuang *et al*., 2019). Therefore, the connection of Lrs1 to the Las and Rhl QS systems was investigated in the strain PAO1-L. Lrs1 overexpression did not affect the transcriptional activity of the *lasR, lasI* and *rhlI* promoters, neither impacted the levels of their respective QS signal molecules (Supplementary Figure 5). Furthermore, the LasR regulated virulence factor elastase (LasB) was not affected by Lrs1 (Supplementary Figure 5 D). Therefore, Lrs1 does not seem to be involved in the regulation of Las or Rhl QS systems in the strain PAO1-L under the tested conditions.

### Lrs1 is involved in the regulation of iron acquisition systems in PAO1-L

To unravel the role of this sRNA in PAO1-L, transcriptomic analysis was performed in the strain overexpressing *lrs1* in *trans* and compared to that of the WT carrying the empty vector. The *lrs1* deletion mutant was not included here to avoid any confounding effects from reduced transcription of the *pqsABCDE* operon. The two strains were cultured in LB in the presence of L-arabinose for the induction of *lrs1*, and samples were collected from both exponential and stationary phase for analysis. (Supplementary Figure 6 A).

The transcriptomic analysis revealed a strong connection between Lrs1 and iron acquisition genes (Figure 2 A, B). Of the 58 upregulated genes in stationary phase, 43 were categorised by gene ontology enrichment analysis in connection to iron acquisition and transport, which was also confirmed by comparing the present dataset to a previous transcriptomic analysis on iron depletion (Ochsner *et al*., 2002) (Supplementary Figure 6 B, C, Supplementary Table 1).

**Figure 2.**
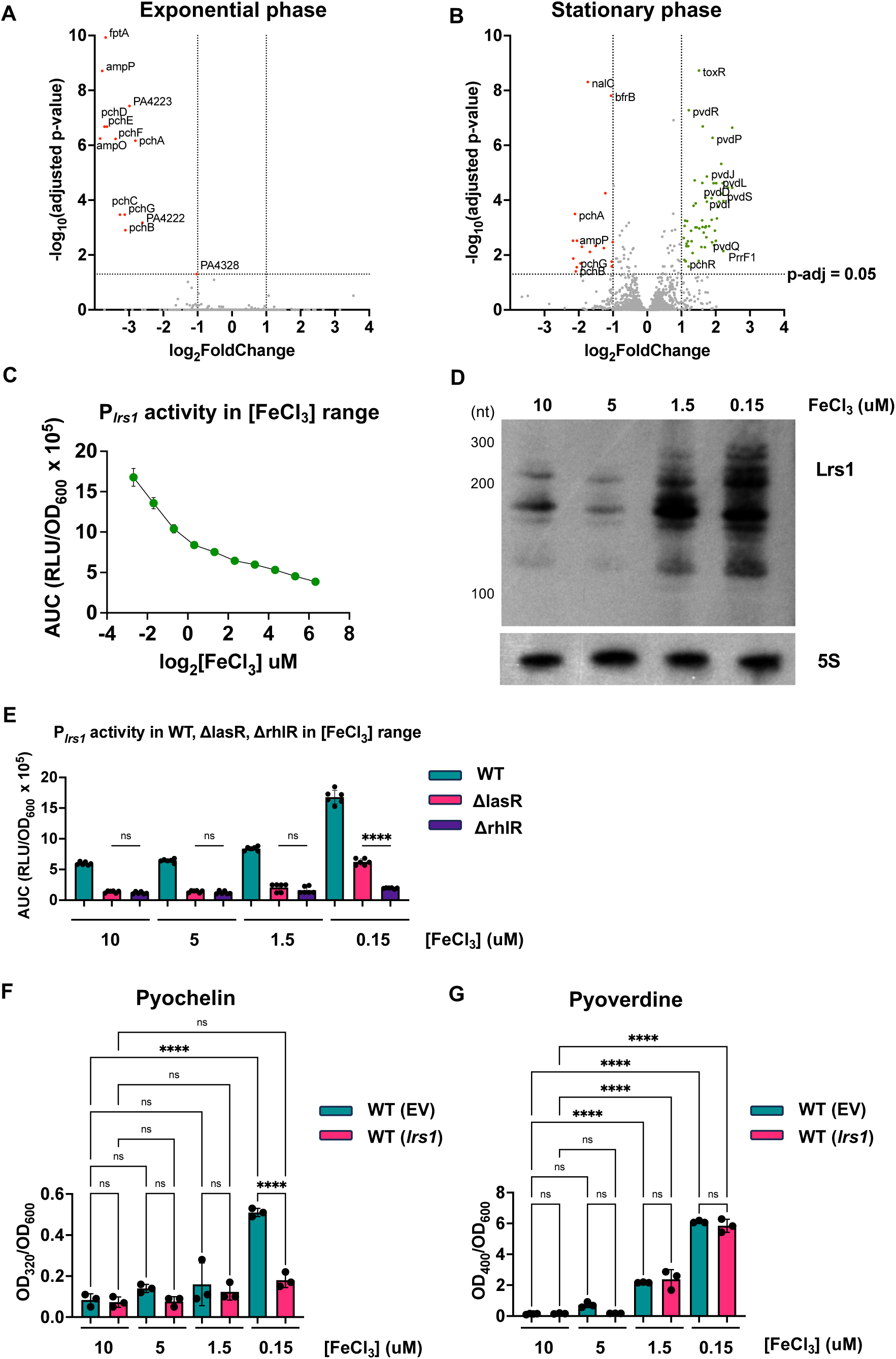
Lrs1 participates in regulation of iron uptake through differential regulation of the siderophores pyochelin and pyoverdine. Volcano plots of the differentially regulated genes in PAO1-L overexpressing *lrs1* at A) exponential and B) stationary phase in LB. Genes with at least −1 ≥ log2FoldChange or log2FoldChange ≥ 1 and p-adjusted ≤ 0.05 were considered differentially regulated. Negatively affected genes are coloured red and positively affected are green. C) Promoter activity of P*_lrs1_* in range of FeCl_3_ concentrations. WT with the P*_lrs1_* bioluminescent reporter was monitored for 48 h in M9 minimal medium with succinate as carbon source. The promoter activity was measured as Relative Luminescent Units normalized by growth (RLU/OD_600_) and then the area under the curve (AUC) was plotted here. The FeCl_3_ concentrations ranged from 80 uM to 0.155 uM following 2-fold serial dilution steps. Error bars represent standard deviation of three biological replicates with two technical replicates. D) Northern blot of Lrs1 in range of iron concentrations. PAO1-L was grown in M9 minimal medium with glucose and 10, 5, 1.5, 0.15 uM FeCl_3_ for 16 h when RNA samples were collected. 5S rRNA was used as a loading control. E) *Lrs1* promoter activity (P*_lrs1_*) in ΔlasR or ΔrhlR compared to PAO1-L WT at 10, 5, 1,5, and 0.15 uM FeCl*_3._* The activity of the P*_lrs1_* bioluminescent reporter was measured throughout growth for 48 h and the Area Under the Curve (AUC) was plotted here. Error bars represent standard deviation of three biological replicates with two technical replicates. F) Pyochelin and G) Pyoverdine quantification in culture supernatants in presence of 0.2% L-arabinose in M9 minimal medium with glucose. Error bars represent standard deviation of three biological replicates. Two-tail ANOVA was used as statistical test. ns: not significant. * : p-value ≤ 0.05, ** : p-value ≤ 0.01, *** : p-value ≤ 0.001, **** : p-value ≤ 0.0001

Among the differentially regulated genes, seven out of the total 14 extracytoplasmic functioning sigma factors (ECFσ) participating in the iron stringency response were identified, displaying higher transcript levels when *lrs1* was overexpressed. Besides the ECFσ factors, their collective regulome was also affected by Lrs1. More than half of the positively differentially regulated genes (35/58) belong to the regulomes of one of these ECFσ factors (Supplementary Table 1 ). Lrs1 may not directly affect all these genes, but its overall impact leads to the activation of the iron acquisition response.

### *Lrs1* is highly transcribed in low iron conditions and this expression is dependent on *RhlR* and *LasR*

To corroborate the importance of Lrs1 in iron stringency, the promoter activity of *lrs1* was measured using a range of iron concentrations in the medium. The activity of P*_lrs1_* was inversely correlated to the iron concentration, showing a strong regulation of Lrs1 by iron levels (Figure 2 C). Northern blot analysis confirmed that Lrs1 transcript levels were higher in lower iron concentrations (Figure 2 D, Supplementary Figure 6 F). The impact of iron depletion on *lrs1* transcription was also dependent on the carbon source used in minimal medium, since Lrs1 was more abundant when the less preferred carbon source glucose was used in place of the more preferred succinate (Figure 2 D, Supplementary Figure 6 F).

The activity of the *lrs1* promoter was also studied in absence of *rhlR* or *lasR* considering their regulatory connection (Figure 1 E). The promoter activity was dependent upon the presence of both regulators in low iron conditions, since deletion of these genes noticeably reduced *lrs1* transcription, although the *lasR* deletion in the lowest iron concentration was less impactful than the *rhlR* deletion (Figure 2 E). Additionally, the dependency of the *lrs1* promoter on the iron regulator Fur was explored and compared to the Fur-regulated promoter of PrrF1 sRNA (P*_prrF1_*) (Wilderman *et al*., 2004). Unlike P*_prrF1_*, the activity levels of P*_lrs1_* were not affected by the addition of the iron chelator 2,2’-Dipyridyl (DIP) in the medium indicating that this promoter is not under the regulation of Fur (Supplementary Figure 6 G). Taken together, unlike the iron-responsive PrrF1,2 sRNAs, Lrs1 remain QS-regulated in low iron conditions, with RhlR and LasR being the main transcriptional regulators.

### *Lrs1* decouples the production of the siderophores pyochelin and pyoverdine

In both exponential and stationary phases, *lrs1* induction downregulated the pyochelin biosynthetic genes *pchDCBA* and *pchEFGHI*, one of the two siderophores produced by *P. aeruginosa* (Figure 2 A, B). However, the biosynthetic gene clusters for the second siderophore, pyoverdine, were upregulated in stationary phase when *lrs1* was overexpressed. In previous studies, the biosynthesis of the two siderophores followed the same trend, being both upregulated or downregulated in response to iron repleted or depleted conditions (Ochsner *et al*., 2002; Palma, Worgall and Quadri, 2003). Here, Lrs1 seems to be involved in decoupling of the expression of the two siderophores.

To investigate the involvement of Lrs1 in the regulation of the two siderophores, the levels of pyochelin and pyoverdine were measured in four different iron concentrations in M9 medium: 10, 5, 1.5 and 0.15 uM FeCl_3_. These concentrations were chosen as high, intermediate, low, and very low iron conditions. Overexpression of Lrs1 reduced pyochelin levels only at the lowest iron concentrations without affecting the levels of pyoverdine, confirming the differential regulation of the two siderophores by Lrs1, a phenomenon not dependent on the carbon source used (Figure 2 F, G, Supplementary Figure 6 I, J). There is a possibility that the increase in iron uptake genes and pyoverdine biosynthesis in the RNA-seq was in response to reduced pyochelin production during the exponential phase of growth. That may have been due to reduced influx of iron from ferripyochelin perceived by the cells as lack of iron resulting in the expression of iron uptake genes.

### *Lrs1* may be involved in other metabolic pathways

RNA-seq uncovered a regulatory connection of between Lrs1 and iron acquisition. Fur would have been a probable target of Lrs1, however, only a specific subset of the Fur regulome was affected by Lrs1 and the siderophore dysregulation did not agree with this hypothesis (Cornelis, Matthijs and Van Oeffelen, 2009). Considering the downregulation of the pyochelin gene clusters in both exponential and stationary phase, it was thought that Lrs1 may impact the expression of the pyochelin transcriptional inducer PchR (Michel *et al*., 2005). However, *pchR* transcription was unaffected in exponential phase and was upregulated in stationary phase (Figure 2 A, B, Supplementary Table 1), hence it appears less likely that Lrs1 is involved in the regulation of *pchR*.

To explore which RNAs may be directly targeted by Lrs1, GRIL-seq (Global sRNA Target Identification by Ligation and Sequencing) was employed (Han, Tjaden and Lory, 2016). Using GRIL-seq, RNAs in direct contact or in proximity to the sRNA of interest can be identified pointing towards direct RNA targets. The RNAs in contact with the sRNA are ligated to the sRNA by an heterologously expressed T4 RNA ligase creating chimeric RNAs. In this study, samples of two biological replicates, from both exponential and stationary phase were collected and searched for Lrs1 chimeras. The chimeras identified in both biological replicates were considered as more likely targets of Lrs1.

None of the pyochelin genes were identified as direct targets of Lrs1 (Supplementary tables 2, 3, 4). On the contrary, the products of the identified mRNAs participate in metabolic pathways including virulence factor production and regulation, carbon metabolism, and adaptation to low oxygen condition, among others. One of the identified potential targets was the *rsaL* mRNA. Although no impact on the *las* QS system by Lrs1 was observed in PAO1-L (Supplementary Figure 5), *rsaL* may have been the regulatory link between *lrs1* and *lasR* in PA14 (Wurtzel *et al*., 2012; Chuang *et al*., 2019).

In addition to the mRNAs, Lrs1 was also found to bind to three sRNAs, CrcZ, PhrS, and ErsA. It has been previously shown that at least PhrS interacts with both CrcZ and ErsA, and that CrcZ interacts with Prrf1 *in vivo* (Han, Tjaden and Lory, 2016; Zhang *et al*., 2017; Gebhardt *et al*., 2023). These RNAs potentially interact directly with Lrs1 and are not necessarily impacted in LB. It is likely that Lrs1 indirectly affects pyochelin biosynthesis through these targets.

Comparison of the potential targets identified with GRIL-seq to the differentially regulated genes from the RNA-seq resulted in *bfrB* being the only commonly identified gene (Supplementary Figure 7 A, B). BfrB is the major bacterioferritin of *P. aeruginosa* and is responsible for iron storage (Eshelman *et al*., 2017; Rivera, 2017). Lrs1 interacted with the 5’ UTR of *bfrB*, near its predicted RBS. Although the two RNAs interacted *in vitro,* translational repression of *bfrB* by Lrs1 could not be verified (Supplementary Figure 7 C). It is possible that either Lrs1 does not majorly impact *bfrB*, or other regulators are masking its effect.

sRNA-mRNA interactions initiate at a short regions of perfect complementarity between the sRNA and the mRNA termed seed region (Wagner and Romby, 2015). Mapping of the seed regions on the mRNA that interacted with Lrs1 revealed that Lrs1 does not only interact with the 5’ UTR of its targets, but also with other regions and with a preference for the 3’ UTR. Lrs1 may not only interact with a diverse number of mRNAs, but the mechanism of action may vary for each mRNA target. There are only a few instances where sRNAs also interact with other regions of their targets besides the 5’ UTR, as was the case of Prrf1 and sKatA (Han, Tjaden and Lory, 2016; Jørgensen, Pettersen and Kallipolitis, 2020).

### *Lrs1* overexpression favours amino acid biosynthesis over pyochelin and alkylquinolones in glucose M9 medium

To further confirm the involvement of Lrs1 in the regulation of pyochelin production, RT-qPCRs were conducted between the WT with the empty vector and WT overexpressing *lrs1* in 0.15 µM iron in M9 with glucose. Unexpectedly, there was no change in the mRNA levels of the pyochelin biosynthetic genes between the two strains (Figure 3 A, Supplementary Figure 9 D). Probably, derepression of the *pch* genes by Fur in low iron has a more dominant effect on their transcript levels than Lrs1 in these conditions.

**Figure 3.**
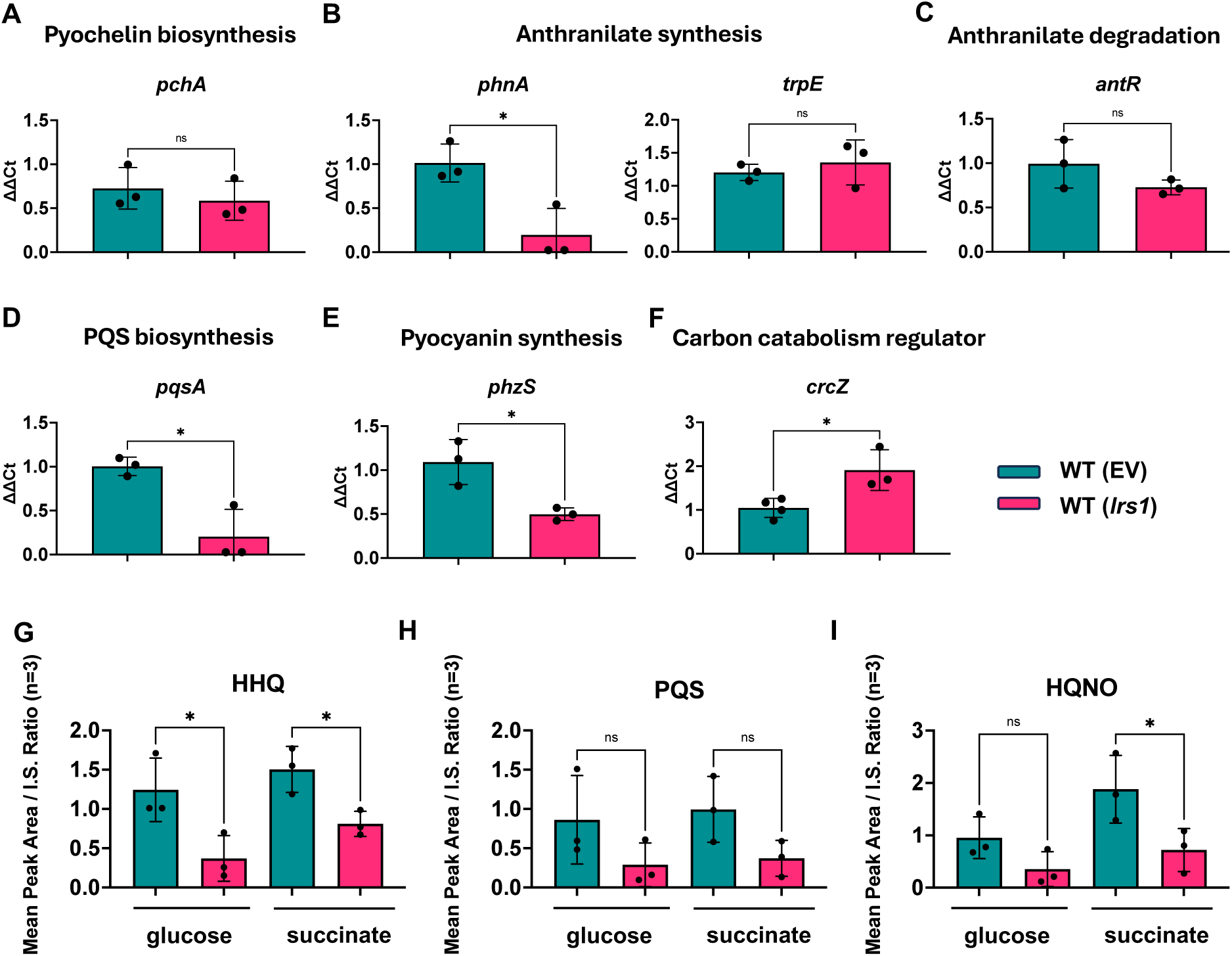
Lrs1 may participate in metabolic rewiring in low iron conditions. RT-qPCR analysis of genes participating in A) pyochelin biosynthesis, B) Anthranilate synthesis, C) Anthranilate degradation, D) PQS biosynthesis, E) pyocyanin production, F) Carbon catabolism regulation in the WT and the *lrs1* overexpressing strain growing in M9 minimal medium with glucose and in the presence of 0.15 uM FeCl_3_. G) HHQ, H) PQS, I) HQNO quantification in 0.15 uM FeCl_3_ in presence of either glucose or succinate. Cells were grown for 20 h, at which point samples were collected. Error bars represent standard deviation of three biological replicates. T-test was used for statistical analysis. ns: not significant. * : p-value ≤ 0.05, ** : p-value ≤ 0.01.

Considering that Lrs1 was ligated to multiple genes participating in carbon catabolism in the GRIL-seq experiments, among them the carbon catabolism regulator CrcZ, it is possible that a metabolic shift might reduce pyochelin precursors and hence production of the siderophore. The precursor of pyochelin is chorismate, which can also be used for the biosynthesis of pyocyanin, and in the biosynthesis of the aromatic amino acids phenylalanine, tyrosine and tryptophan, or its precursor, anthranilate. Anthranilate can be used for the biosynthesis of alkylquinolones, or directed to the TCA cycle. To investigate these possibilities, RT-qPCR analysis was conducted on genes of these metabolic pathways, to see if they were affected by overexpressing *lrs1*. Indeed, the levels of mRNA of genes for anthranilate synthase *phnAB*, *pqsA*, and the pyocyanin biosynthetic genes were all downregulated in the *lrs1* overexpressing strain (Figure 3 B, D, E, Supplementary Figure 9 A, C). The gene *phzS*, which was identified as another potential direct target of Lrs1, was also downregulated, a phenomenon that could be partly through direct interaction with Lrs1. However, genes of anthranilate degradation (*antA*) or tryptophan biosynthesis, *trpEG*, were unaffected (Figure 3 B, C, Supplementary Figure 9 A, B). Although anthranilate for alkylquinolones synthesis can be supplied by either the PhnAB or the TrpEG anthranilate synthases, the primary destination of anthranilate produced by TrpEG is tryptophan biosynthesis (Palmer, Jorth and Whiteley, 2013). The carbon catabolism regulatory sRNA CrcZ, another potential target of Lrs1, was upregulated in the *lrs1* overexpressing strain, in line with the hypothesis of Lrs1 involvement in metabolic rewiring (Figure 3 F). To corroborate these findings, alkylquinolones were quantified in low iron conditions. Indeed, *lrs1* overexpression significantly reduced the levels of HHQ but did not impact PQS levels, and HQNO was only impacted when the carbon source was succinate (Figure 3 G, H, I). These results indicate that shared metabolites of these pathways are redirected towards either the TCA cycle or aromatic amino acid biosynthesis, deprioritising biosynthesis of pyochelin, alkylquinolones, and pyocyanin in low iron conditions.

## Discussion

Post-transcriptional regulation plays crucial role in microbial adaptation to environmental changes. A number of sRNAs have already been implicated in controlling the QS systems of *P. aeruginosa* (Oglesby *et al*., 2008; Sonnleitner *et al*., 2011; Carloni *et al*., 2017; Thomason *et al*., 2019). A previous study has pointed the sRNA Lrs1 in the regulation of the Las QS system in the strain PA14 (Chuang *et al*., 2019). However, no change in the regulation of *lasR* by Lrs1 was identified in PAO1-L in the current study. It is noted that this regulation was mainly observed in surface-attached cells in PA14, whereas here planktonically grown cells were used. Considering that reduced elastase activity was reported in the *lrs1* mutant in PA14 growing under aerobic conditions in LB and reduced *lasR* transcript levels were observed in the same mutant in planktonic cells, it was reasoned this phenomenon could be observed here too. GRIL-seq indicated *rsaL* mRNA as a candidate target of Lrs1 although RNA-seq did not reveal changes in the relative abundance of the *rsaL* mRNA when *lrs1* was overexpressed in PAO1-L. It is likely that other regulators may mask the effect of Lrs1 in this strain, but *rsaL* may in fact be the missing link in the regulation of *lasR* by Lrs1 in PA14 (Chuang *et al*., 2019). The strains PAO1 and PA14 are genetically and phenotypically distinct, with regulatory differences between them having been reported before (Grace *et al*., 2022). This might be the case here too, where Lrs1 has adopted separate roles in the two reference strains, adding to the phenotypic diversity within the species. That would not be the first time where an sRNA does not regulate the same targets in the two strains. The sRNA ErsA was reported to downregulate *algC* in PAO1 only, whereas no regulation could be observed in PA14 (Ferrara *et al*., 2015). Hence, sRNAs can be drivers of phenotypic and regulatory differentiation between the two strains, and that might expand within the whole *P. aeruginosa* species.

Iron stringency in *P. aeruginosa* is tightly regulated for optimal acquisition and utilisation of this metal. *P. aeruginosa* produces the siderophores pyochelin and pyoverdine for iron uptake, altering between the two depending on the relative abundance of iron in the medium (Dumas, Ross-Gillespie and Kümmerli, 2013). Pyoverdine and pyochelin differ in their affinity for iron with pyoverdine having a higher Kd than pyochelin making it more useful in iron scarcity (Brandel *et al*., 2012) Here, the sRNA Lrs1 may participate in the regulatory switch between the two siderophores in PAO1. Only pyochelin levels were reduced in the lowest iron concentration tested here, with pyoverdine being unaffected in the *lrs1* overexpressing strains. So far, most studies have showed parallel increase or decrease in the expression of the siderophore biosynthetic operons (Ochsner *et al*., 2002; Wilderman *et al*., 2004; Nelson *et al*., 2019). The only exception has been the sRNA PrrH, which is involved in pyochelin regulation, with the effect being altered in presence of light or growing the cells statically or aerobically (Hoang *et al*., 2023). Here, Lrs1 may participate in the differential regulation of the siderophores based on the nutritional needs of the cells. Along with the sRNAs Prrf1, Prrf2, and PrrH, Lrs1 may be involved in the iron stringency response of *P. aeruginosa* by decoupling the synthesis of the two siderophores. By doing so, valuable resources are directed towards essential processes, such as pyoverdine and amino acid biosynthesis, increasing the adaptation of the organism and its competitiveness in the environment.

This is also evident by the conditional negative impact of Lrs1 on the Pqs QS system in low iron. PQS can act as an iron scavenger, enhancing the iron uptake response of *P. aeruginosa* (Bredenbruch *et al*., 2006). Previous studies have indicated that pyochelin and PQS may share the same transporters to enter into the cells (Lin *et al*., 2017; Zhang *et al*., 2024). PQS levels remain high under low iron, in part due to the downregulation of the anthranilate degradation transcriptional regulator *antR* by Prrf1,2 and PhrS, providing more precursors for PQS biosynthesis (Djapgne *et al*., 2018; Dubern *et al*., 2023; Gebhardt *et al*., 2023). Lrs1 may provide a regulatory link between pyochelin and PQS, further connecting the PQS QS system to iron uptake. Here, it is proposed that the RhlR-regulated sRNA Lrs1 amplifies the effects of RhlR and CrcZ on the fates of chorismate and anthranilate (Figure 4). Lrs1 increases the

**Figure 4.**
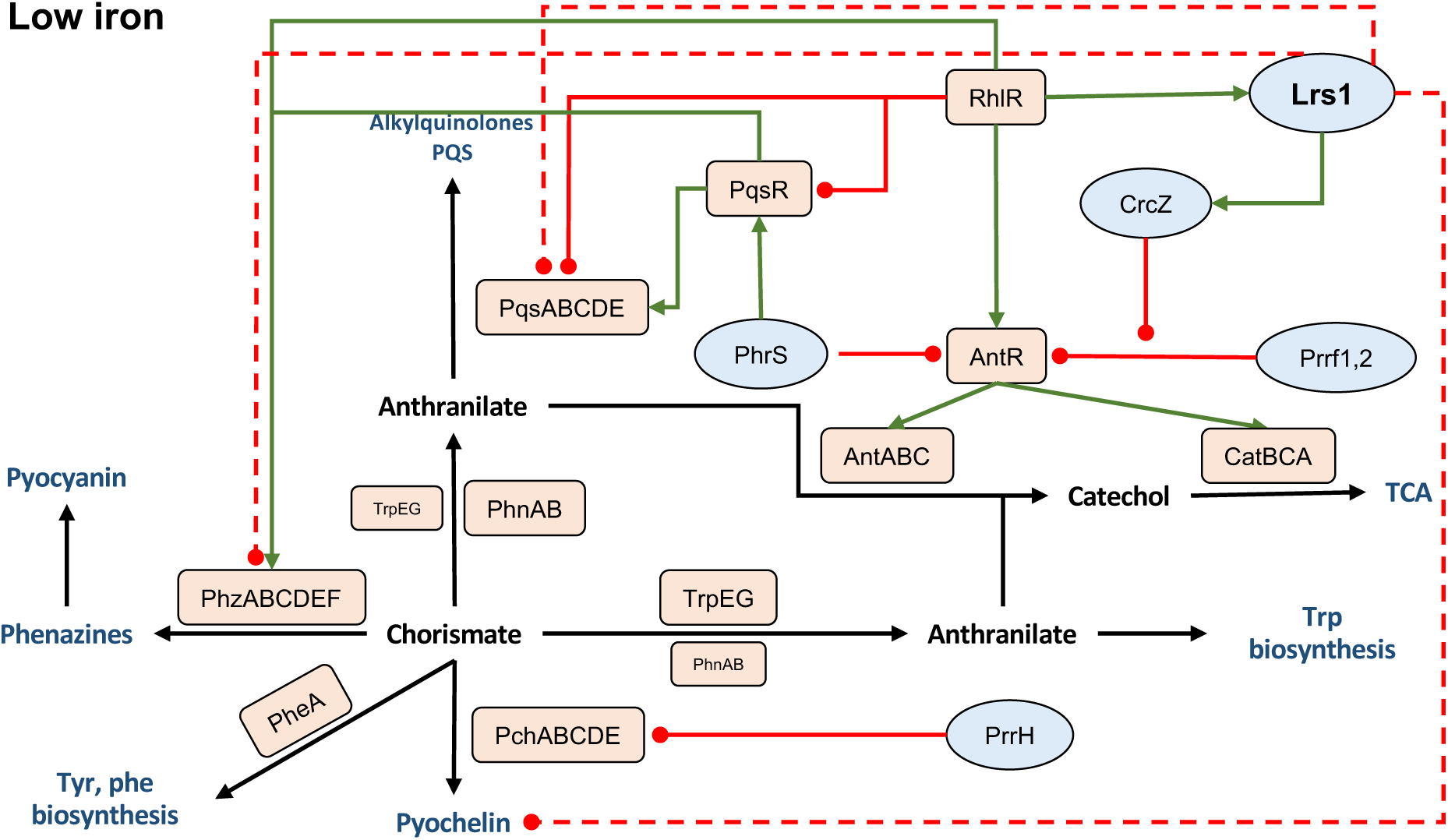
Proposed model of Lrs1 involvement in balancing metabolic flow in low iron in *P. aeruginosa* PAO1-L. Chorismate is the precursor of the aromatic amino acids, tyrosine, and phenylalanine, as well as pyochelin, pyocyanin, and anthranilate. In turn, anthranilate can be used for the synthesis of the aromatic amino acid tryptophan, the production of alkylquinolones and the QS molecule PQS, or it can be catabolized via the TCA cycle. In low iron, the low-iron affinity siderophore pyochelin is expressed for iron scavenging, along with increased production of PQS that is used both for QS activation and iron entrapment. The transcription of the PQS biosynthetic genes *pqsABCDE* is activated by the Pqs transcriptional regulator PqsR and indirectly by the PhrS sRNA. At the same time, PhrS downregulates the gene of the anthranilate degradation transcriptional regulator AntR and by extension the anthranilate degradation genes *antABC* and *catBCA* preserving anthranilate for PQS production. The iron stringency response sRNAs Prrf1,2 further repress *antR* sparing precursor molecules for PQS production synthesis while PrrH represses pyochelin biosynthesis. The sRNA CrcZ interferes with the Prrf1,2 mediated repression of *antR,* promoting the degradation of anthranilate for energy conversion. The Rhl transcriptional regulator RhlR further promotes anthranilate degradation by downregulating *pqsR* and *pqsABCDE* while upregulating *antR*. RhlR, along with LasR, upregulate Lrs1 which enhances the effect of RhlR and CrcZ. Lrs1 negatively impacts pyochelin, preserving chorismate for aromatic amino acid biosynthesis. Additionally, Lrs1 downregulates alkylquinolone synthesis through both the *pqs* operon and the anthranilate synthase genes *phnAB* without affecting the anthranilate synthase genes *trpEG* redirecting anthranilate towards either degradation via the TCA cycle or to tryptophan biosynthesis. Finally, Lrs1 increases CrcZ levels, further promoting metabolic efflux to amino acid biosynthesis and energy conversion. Proteins are depicted in orange rectangles and sRNAs in blue ovals.

levels of CrcZ possibly through direct interaction although the mechanism of action has not been fully elucidated. The sRNA CrcZ interferes with the Prrf1,2-mediated regulation of *antR* with a possible negative effect on Pqs (Sonnleitner, Prindl and Bläsi, 2017). Additionally, RhlR negatively affects the Pqs system by repressing *pqsR* and upregulating *antR* (Xiao, He and Rahme, 2006; Oglesby *et al*., 2008). The pool of available anthranilate for alkylquinolones is reduced as more of it is supplied into the TCA cycle, through the actions of the AntABC enzymes. At the same time, the anthranilate synthase genes *phnAB* supplying anthranilate for PQS production are also downregulated. However, the *trpEG* anthranilate synthase genes are unaffected due to their central role in tryptophan biosynthesis (Palmer, Jorth and Whiteley, 2013). Phenazine and pyochelin synthesis is reduced, sparing more chorismate for amino acid biosynthesis. Taken together, Lrs1 may participate in balancing metabolic flow, favouring essential processes such as amino acid biosynthesis and energy conversion in low iron conditions.

*In vivo* interaction of Lrs1 with multiple mRNAs and sRNAs implicated in diverse metabolic pathways without a clear connection to iron, suggests that Lrs1 may have a more versatile regulatory role. The dysregulation of these pathways was not apparent in the RNA-seq carried out in the present study, which indicates that regulation may manifest in environmental conditions other than LB. That was the case of CrcZ upregulation, observed here in low iron conditions only. Another hypothesis is that Lrs1 may not act through altering transcript levels of these genes. In many of the targets identified in the GRIL-seq experiment, Lrs1 was bound to their 3’ UTR, suggesting a distinct function. In a recent study, a similar pattern was observed where sRNAs were bound to the 3’ UTR of mRNAs in *P. aeruginosa,* among them the 3’ UTR of *oprB* (Gebhardt *et al*., 2023). It was hypothesised that an sRNA may be transcribed from the 3’ UTR consisting the actual sRNA target, similarly to the 3’ UTR derived sRNA sKatA interacting with PrrF1 and PrrF2 (Han, Tjaden and Lory, 2016). Here, Lrs1 was also bound to the 3’ UTR of *oprB*, strengthening the possibility of the existence of this sRNA. Collectively, these interactions may indicate a complex regulatory network where sRNAs interact with their targets but also existing in an equilibrium with the other sRNAs and 3’ UTR-derived sRNAs. The 3’ UTR-derived sRNA-sRNA base-pairing could function as a titrating mechanism to prevent the sRNAs acting upon their targets. That offers a way to controlling the concentration of available sRNA to base-pair with its targets in the presence of competing sRNAs.

This study highlights the importance of sRNAs for survival of *P. aeruginosa* under nutrient starvation. Here, metabolic fate in low iron seems to be determined by five sRNAs: Lrs1, CrcZ, PhrS, and Prrf1,2 acting in opposite ways to each other. The critical role of sRNAs in low iron may be a way for *P. aeruginosa* to navigate an ever-fluctuating environment where nutrient availability can be stochastic. The sRNAs quickly alter the RNA landscape, favouring the more beneficial metabolic pathways for the current conditions, tipping the balance in favour of *P. aeruginosa* on interspecies competition and infection establishment.

## Acknowledgements

This project was funded by the Wellcome Trust Antimicrobials and Antimicrobial Resistance DTP programme, [Award 108876/Z/15/Z], the National Biofilms Innovation Centre (NBIC), which is an Innovation and Knowledge Centre funded by BBSRC, Innovate UK and Hartree Centre [Awards BB/R012415/1 and BB/X002950/1], and ANID/Becas Chile, Doctorado en el extranjero [file number 72710364]. We would also like to thank Professor Stephen Lory from Harvard Medical School, for kindly providing the vector system for the GRIL-seq experiment.

## Experimental Procedures

### Strains, plasmids, primers, and culture media

The bacterial strains and plasmids used in this study are listed in Supplementary Tables 5 and 6. *P. aeruginosa* strain PAO1-L was used throughout this study unless otherwise stated. *E. coli* DH5α was used for routine cloning and *E. coli* S17-1 for conjugal transfers. All strains were grown overnight in lysogeny broth (LB) at 37°C with shaking at 200 rpm, or on LB agar plates. LB was prepared according to (Sambrook and Russell, 2001). For iron-free conditions, M9 Basal salts (Basal salt solution 2x concentrate: 95.9 mM Na_2_HPO_4_, 44 mM KH_2_PO_4,_ 34.2 mM NaCl; Supplements 100x concentrate: 1.9 M NH_4_Cl, 20 mM CaCl_2,_ 200 mM MgSO_4_) were deferrated with Chelex 100 (50-100 Mesh, Sigma Aldrich), and the supplements were prepared in iron-free water. As carbon source, iron-free sodium succinate or glucose were added to a final concentration of 20 mM. For low iron conditions, 0.15 uM FeCl_3_ was added to the medium, and for iron-replete conditions, the final ferric iron concentration was 10 uM. Antibiotics were added to the media at the following concentrations when necessary: for *E. coli*, 10 ug/mL tetracycline (Tc), 10 ug/mL gentamicin (Gm), 100 ug/mL ampicillin (Ap); for *P. aeruginosa*, 125 ug/mL Tc, 20 ug/mL Gm, 200 ug/mL carbenicillin (Cb). Isopropyl β-D-1-thiogalactopyranoside (IPTG) to a 1 mM final concentration was added when required at the start of the culture. Blue-white selection of colonies was done by adding 5-bromo-4-chloro-3-indolyl-β-D-galactopyranoside (x-Gal) at 40 uM final concentration in agar plates. For the overexpression of *lrs1* from the pKH6 vector, L-arabinose was added in the medium to a final concentration of 0.2% V/V. Oligonucleotides used in this study are listed in Supplementary Table 7.

### Vector construction

The plasmid pBS_1 was constructed by PCR amplification of a 972 bp region with primers LRS1RV and LRS1FW, which included 615 bp upstream of *pqsA* and 357 bp downstream of the *pqsA* translational start site, and after digestion, it was ligated between the PstI and EcoRI sites of pBluescript.

The plasmid pDP013 was constructed by PCR amplification of the upstream and downstream region of the *lrs1* gene with primer pairs lrs1190delF1FW and lrs1148delF1RV, lrs1190delF2RV and lrs1148delF2FW, respectively. The PCR products were then cloned into PstI digested pTS1 with the HiFi DNA assembly kit (NEB). Only the first 138 bp of the *lrs1* gene were deleted, excluding the TSS2, in order to avoid deleting the rpoN binding site situated directly downstream and has been shown to control *pqsA* expression (Shao *et al*., 2018).

The plasmid pKH6-lrs1 was constructed by cloning *lrs1* in the vector pKH6. The vector pKH6 contains the arabinose-inducible P*_bad_* promoter and has been designed for overexpression of sRNAs (Han *et al*., 2016). The transcriptional start site of P*_bad_* is an

adenosine, and is located 2bp downstream of an XbaI restriction site (Han *et al.,* 2016). The sRNA gene was amplified with the primers LRS1XBAIFW and LRS1X_ECORI_RV, and was subsequently ligated between the XbaI and EcoRI sites of pKH6. An adenosine was added directly upstream of their transcriptional start site, in order to allow for correct transcriptional initiation from the P*_bad_* promoter. The length of cloned *lrs1* was 248 bp as this was previously used in (Chuang *et al*., 2019).

For the transcriptional promoter fusions in pminiCTXlux, promoter regions were cloned between the HindIII and PstI sites in the MCS of pminiCTXlux. P*_lrs1_* was amplified with primers LRS1PROHINDIIIF and LRS1PROPSTIRV, P*_lasR_* with PlasRHindIIIFW and PlasRPSTIRV, P*_lasI_* with PlasIHindIIIFW and PlasIPSTIRV, and P*_rhlI_* with PrhlIHindIIIFW and PrhlIPSTIRV. All promoters were amplified from the PAO1-L gDNA. P*_pqsA_* was amplified from pBS_1. In the case of translational promoter fusions to pminiCTXlux, DNA fragments containing the promoters were cloned in the XcmI restriction site directly upstream from the translational start site of *luxC* on pminiCTXlux by HiFi DNA assembly (NEB) following the manufacturer’s instructions. The promoter P*_pqsA_* was amplified using the primers PpqsA-translational-F and PpqsA-translational-R from plasmids pBS_1.

The pBx-rne1 vector was constructed with Golden Gate assembly. Briefly, the single stranded oligonucleotides sgRNA_1_RV and sgRNA_1_FW were dimerised by incubating equimolar concentration of each oligonucleotide in annealing buffer (10 mM Tris, pH 8.0, 50 mM NaCl, 1 mM EDTA). The assay was heated to 95°C for 5 min and then the temperature slowly decreased to 25°C. The double stranded oligo was then incubated with the vector pBx-Spas-sgRNA-Gm and BbsI restriction digestion enzyme in rCutsmart buffer (NEB) for 10 min at 37°C. Afterwards T4 DNA ligase and 10 mM ATP were added to the assay. The reaction was incubated for 20 circles of alternate steps, 3 min 37°C and 3 min at 16°C.

The plasmids pME6032-*lasR* and pME6032-*rhlR* were constructed by cloning the open reading frames of *lasR* and *rhlR* between the EcoRI and KpnI sites, *in frame* with the translational initiation start site at the end of the P*_tac_* promoter in pME6032. The gene of *lasR* was amplified with primers LasR-C-F and LasR-C-R, *rhlR* with RhlR-C-F and RhlR-C-R from the genome of PAO1-L.

The suicide vectors pME3087-rhlR, pME3087-rhlI, pME3087-lasR, and pME3087-lasI for the construction of deletion mutants of the genes *rhlR, rhlI, lasR, and lasI,* respectively, were constructed as follows. The flanking regions of each gene were amplified with their respective primers and were individually cloned in pBluescript. Then, the upstream and downstream regions were cloned and ligated together in pBluescript at their mutual XbaI site, or BamHI for the pME3087-pqsE, with restriction digestion and ligation. The resulting fragment was transferred to the suicide vector pME3087 that was then used for the construction of the respective deletion mutants.

### Strain construction

All vectors were transferred to *E. coli* electrocompetent cells prepared according to Sambrook and Russel (Sambrook and Russell, 2001). *P. aeruginosa* electrocompetent cells were prepared according to (Choi, *et al*., 2006).

For deletion mutants, the corresponding suicide plasmids were transformed in S17-1 and subsequently conjugated to PAO1-L. Following conjugation, strains carrying a pTS1 or pEX18 based suicide vector were sucrose counter-selected for double recombinants according to the protocol from (Huang and Wilks, 2017). Colonies resistant to sucrose and susceptible to the corresponding antibiotic were PCR screened to identify the mutants followed by sequencing to confirm the deletion. Forthe construction of deletion mutants with pME3087 based suicide vectors, the tetracycline enrichment approach was followed. Briefly, cells from an overnight culture were incubated in LB for 2 h, 10 ug/mL tetracycline was added for 1 h followed by addition of 300 ug/mL carbenicillin for 6 h. This process was repeated three times after which tetracycline sensitive colonies were selected and screened by PCR. The pminiCTXlux suicide plasmids carrying the promoter fusions were conjugated to *P. aeruginosa* from *E. coli* S17-1.

The vector pUC18-miniTN7-Plac-dCas9-Gm was transferred to PAO1-L by triparental conjugation with the vector pTNS1. Following insertion of the vector to the chromosome, the gentamycin resistance cassette was excised by electroporating the flipase encoding vector pFLP2. Curation of pFLP2 was done by sucrose counterselection as described in (Huang and Wilks, 2017). Afterwards, pBx-Spas-sgRNA-Gm and pBx-rne1 were electroporated to the gentamycin sensitive strain.

### Bioluminescence assays

Promoter activity was determined using pminiCTXlux bioluminescent reporters integrated to the *P. aeruginosa* chromosome. Cultures were grown overnight in LB with appropriate antibiotics. The overnight cultures were diluted to a final OD_600_ = 0.01 and 200 uL were added in each well in a 96-well plate (flat black, transparent bottom, GreinerBioOne), in triplicates, and the plate was covered with a transparent lid. Measurements of OD_600_ and relative luminescence units (RLU) were taken every 30 minutes for 24 h at 37°C unless otherwise stated in the figure legend. Promoter activity was calculated as RLU/OD_600_. Statistical analysis was performed with the software Graphpad Prism.

### Pyoverdine and pyochelin quantification

In 100 mL flasks, 10 mL iron-free M9 minimal media with 20 mM glucose was inoculated with 0.05 %V/V overnight cultures and was incubated for 20 h at 37 °C, 200 rpm. Following centrifugation to pellet the cells, the supernatants were filter sterilised. Pyoverdine was quantified by measuring absorbance of the supernatant at 400 nm. Pyochelin was quantified according to (Cunrath *et al*., 2020). Briefly, 1 mL of supernatant was acidified with 50 uL 1 M citric acid. Then, pyochelin was extracted twice with 500 uL ethyl acetate and absorbance was measured in a quartz cuvette at 320 nm. Both OD_400_ and OD_320_ were divided by OD_600_ to account for cell culture density.

### Elastase quantification

In 50 mL flasks, 10 mL LB was inoculated with 0.05% V/V overnight cultures and was incubated for 20 h at 37 °C, 200 rpm. 1.5 mL of culture was centrifuged to pellet the cells, and the supernatant was filter sterilised. 100 uL of the supernatant was mixed with 900 uL ECR buffer (1.41 g/100 mL Tris, 19.47 mg/ 100 mL CaCl_2_, pH = 7.5) with 30 mg/mL elastin congo red in eppendorfs. The mix was incubated at 37 °C, 200 rpm for 4 h. Then, the eppendorfs were centrifuged at 13000 g, 3 minutes, and the absorbance of the supernatant was measured at 495 nm.

### Quantification of AQs and anthranilate

The comparative analysis of alkyl quinolones in supernatant samples was conducted by liquid chromatography – tandem mass spectrometry (LC-MS/MS) using an Excion LC system in tandem with a Sciex Qtrap 6500+ mass spectrometer. Analytical samples were prepared by mixing 10 uL of sterile filtered supernatants with 90 uL of an internal standard solution (500nM solution of deuterated HHQ (d4-HHQ) in MeOH). 5 uL of each sample were injected onto a Phenomenex Gemini C18 column (3.0µm, 50 x 3.0mm), and eluted at an LC flow rate of 0.45 mL/min. A linear binary gradient (mobile phase A: 2 mM 2-picolinic acid in 0.1% (V/V) formic acid, mobile phase B: 0.1% (V/V) formic acid in MeOH) from 10% B to 99% B over 5 min, followed by a further 2 min at this composition was used. AQ detection was conducted with the MS operating in MRM (multiple reaction monitoring) mode, screening the LC eluent for the specific analytes of interest.

### Northern blots

Total RNA was extracted from cell cultures using the RNeasy plus Universal mini kit (Qiagen) following the manufacturer’s instructions. For the detection of Lrs1 by Northern Blot, the DIG (digoxygenin) Northern Starter kit (Roche) was used following the manufacturer’s instructions with some modifications. The endogenous control was 5S rRNA. The probes for both Lrs1 and 5S were created with *in vitro* transcription of PCR templates for both genes. For Lrs1 detection, 10 ug RNA were loaded per sample and 200 ng for 5S rRNA. The samples were loaded in 10.5% V/V vertical denaturing polyacrylamide gels with 5 M urea and 1x TBE. The samples were mixed with equal volume of denaturing RNA loading buffer. As a marker, the Century^TM^-RNA marker templates (Invitrogen) were *in vitro* transcribed and incorporated DIG-11-UTP. T the gels were electroblotted on positively charged nylon membranes (Amersham Hybond-N RPN 203 N) followed by UV crosslinking and hybridised in DIG EasyHyb buffer (Roche) at 68 °C with gentle agitation overnight. Following incubation with anti-DIG antibody, the membranes were incubated with the substrate CDP-star ready-to-use solution (Roche). The membranes were finally developed by exposing to X-ray film.

### Quantitative Real Time PCR (RT-qPCR)

Following RNA extraction, the RNAs were treated with Turbo DNA-free kit to remove residual RNA. The RNA was reverse transcribed with the GoScript Reverse Transcriptase kit with random primers (Promega). The qPCR mix used here was the Power SYBR Green PCR Master Mix (Applied Biosystems, Cat: 4367659). The qPCRs were performed in a QiaQuant 5x multiplex system The *rpoD* mRNA was chosen as endogenous control. Three biological replicates and three technical replicates were analysed for each target. ΔΔCt ratios were calculated with the equation described by (Pfaffl, 2001) and data analysed on GraphPad Prism.

### Electrophoretic Mobility Shift Assay (EMSA)

The RNAs were transcribed *in vitro* with the MegaScript T7 Transcription kit (Invitrogen) from PCR templates. The templates were made for each RNA with a forward primer containing the T7 promoter in the 5’ end. Subsequently, the RNAs were purified with the RNeasy cleanup kit (Qiagen). A non-radioactive approach was followed for the visualisation of EMSA. The tagged RNA and Lrs1 was made with a reverse primer starting with the sequence 5’ TTTTTTTTCCCCCCCCC 3’. This sequence would add a 5’ GGGGGGGGGAAAAAAAA 3’ which was complementary to a probe (the Atto probe) covalently bound to the fluorophore Atto_700_. Atto_700_ fluoresces at the infrared, with peak excitation at 699 nm and peak emission at 716 nm. The sRNA and the Atto-probe were mixed at a 1:5 molar ratio annealing buffer (10 mM Tris-HCl pH=7.5, 50 mM NaCl, 1 mM EDTA). The reaction was boiled for 5 min at 95°C and it was gradually cooled to 25°C. The RNA targets were diluted in water to a 1 uM and boiled for 1 minute at 95 °C, chilled on ice for 1 minute, and transferred at room temperature for 5 minutes to allow for RNA folding (Bak *et al*., 2015). The probed sRNA and the target RNA were mixed together in dimerization buffer (10x: 200 mM Tris-HCl pH = 7.5, 100 mM NaCl, 250 mM MgCl_2_, 10 uM yeast tRNA) in 10 uL reactions. The probed sRNA concentration was constant at 10 nM, and the target RNA was at 0, 10, 25, 50, and 100 nM. The reactions were incubated for 30 min at 37 °C and the reaction was stopped by addition of 2 uL non-denaturing loading buffer (12% glycerol, 1 x TBE, 0.001% w/V bromophenol blue). The whole reaction was loaded in 5% non-denaturing PAGE. The gels were run in 0.5 x TBE, at 50 V, 2 h, at 4 °C, in the dark. The gels were visualised at the Li-cor Odyssey Infrared Imaging System.

### RNA sequencing

RNA samples for RNA-sequencing were harvested as follows. In 50 mL flasks, 10 mL LB with 0.2% L-arabinose was inoculated with 0.05% V/V overnight culture. The cultures were incubated at 37 °C, 200 rpm. Exponential phase samples were taken when cultures reached an OD_600_ ∼ 0.5, and early stationary phase when they reached OD_600_ ∼ 3.3.

The RNA sequencing was conducted in the DeepSeq Facilities at the University of Nottingham. The library pool was sequenced on the Illumina NextSeq 500 using a NextSeq 500 high Output 150 cycle kits (Illumina), to generate over 10 million pairs of 75 bp paired-end reads per sample.

### GRIL-seq

The sample collection and preparation for RNA-Seq was performed as described in the analytical protocol in the supplementary files of (Han *et al*., 2016). In 250 mL flasks, 30 mL LB containing Cb_400_ and Gm_20_ was inoculated with overnight cultures to a final OD_600_ = 0.01. The flasks were incubated at 37 °C, 200 rpm until OD_600_ ∼ 0.5 (exponential phase samples) at which point the cultures were split. 10 mL were added in two 50 mL flasks and 10 uL IPTG 1 M (C_final_ = 1 mM) was added in one flask (test flask, induced conditions) to induce expression of T4 RNA ligase and nothing was added inthe other (control flask, uninduced conditions). An hour later, L-arabinose was added in the test flask to induce the sRNA transcription from the pKH6 vector. 20 minutes later, 1.6 mL samples were collected. The same procedure was followed for the stationary phase samples, with the induction starting when OD_600_ ∼ 3.3. Following sRNA enrichment, the RNA was sequenced at the DeepSeq facilities at the University of Nottingham as described for the RNA-sequencing method above with some modifications. The library pool was sequenced on the Illumina NextSeq500 over one NextSeq500 Mid Output 300 cycle kit (Illumina), to generate approximately 20 million pairs of 150-bp paired-end reads per sample.

Data analysis was conducted on the online server Galaxy (usegalaxy.eu). The reads were locally aligned to the corresponding sRNA sequence only using Bowtie2 (v2.5.0, galaxy version 2.4.5 +galaxy1) (Langmead *et al*., 2009) with the following parameters changed: allowed mismatches in the multiseed alignment: 1, length of the seed substrings: 20, match bonus: The aligned reads were subjected to a further local alignment with Bowtie2 preserving the same parameters. This time the genome for the alignment was that of PAO1 with the sequences of the sRNA and the rRNAs deleted. Mapped reads were counted with FeatureCounts (galaxy version 2.0.3+galaxy1) (Liao, Smyth and Shi, 2014). The mapped alignments were visualised with Artemis (https://www.sanger.ac.uk/tool/artemis/). The prediction of sRNA-RNA seed areas were conducted with the online tool IntaRNA 2.0 (http://rna.informatik.uni-freiburg.de/IntaRNA/Input.jsp) (Busch *et al*., 2008; Mann *et al*., 2017). The alignment of the chimeric reads to a manually made chimeric sequence of the sRNA and the mRNA was done on Benchling (https://benchling.com/) using MAFFT aligner (Katoh and Standley, 2013) with the default parameters.

## Notes

### Competing Interest Statement

The authors have declared no competing interest.

