## Supplementary Figures and Tables for "The Quorum Sensing regulated sRNA Lrs1 is involved in the adaptation to low iron in *Pseudomonas aeruginosa*"

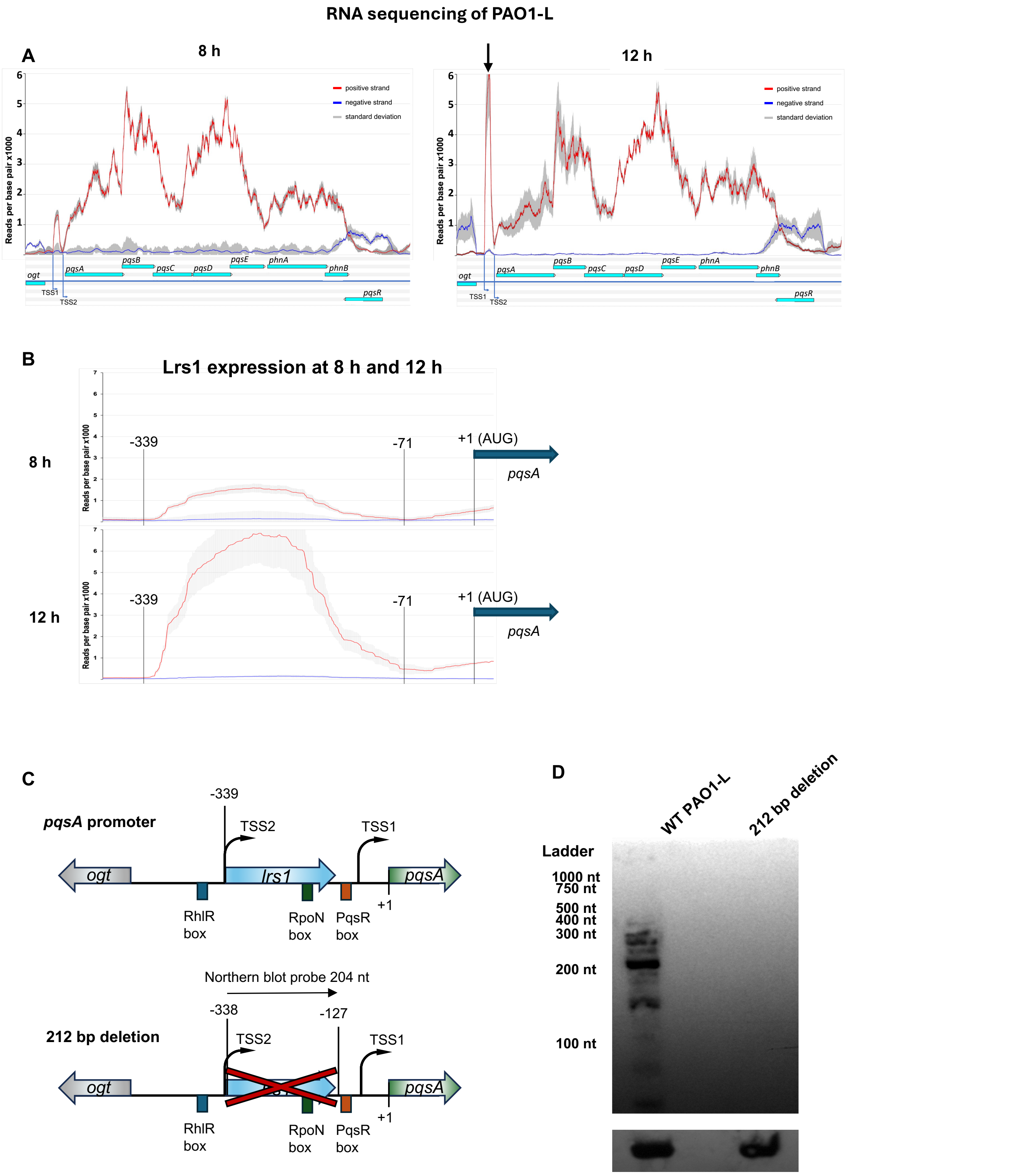


Supplementary Figure 1 Identification of *lrs1* in PAO1-L in RNA-sequencing. PAO1-L was planktonically cultured in LB for 8 and 12 h at which timepoints RNA was extracted for RNA-sequencing. This figure shows read coverage x 1000 at the two chosen timepoints. A) Read coverage of the entire *pqsABCDE* operon, *phnAB* and *pqsR* genes. B) Read coverage of the *pqsA* promoter at 8 and 12 h showing the increased reads between TSS1 and TSS2, the region corresponding to *lrs1*. C) Schematic representation of *lrs1* deletion including 22 bp downstream of the 190 bp end of the gene. D) Northern blot of Lrs1 in the WT and 212 bp deletion strain showing the specificity of the northern blot probe.

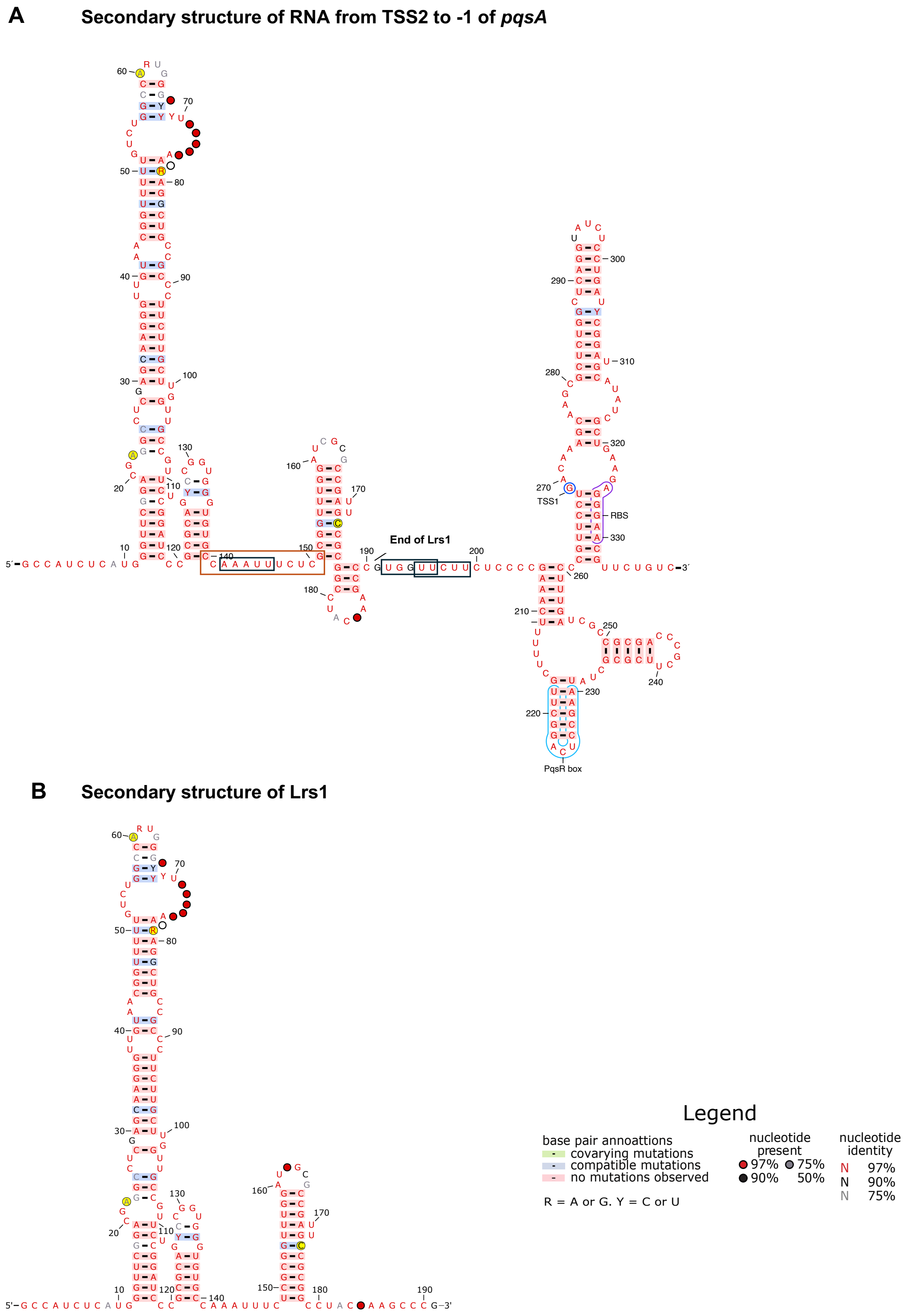


Supplementary Figure 2 Predicted secondary structure of Lrs1 and the area from TSS2 to *pqsA* +1. Predicted secondary of A) TSS2 to *pqsA*, B) 190 nt form of Lrs1. The secondary structures were created by aligning 377 complete *P. aeruginosa* genomes downloaded from Pseudomonas.com, followed by uploading to the online server of RNAalifold (Bernhart *et al.*, 2008) and drawing the structures with R2R (Weinberg and Breaker, 2011). Nucleotides in yellow represent single nucleotide polymorphisms (SNPs) between the PA14 and PAO1 Lrs1 sequences. Sequences in black rectangles are candidate RNase E targets. The RpoN binding site is shown with an orange rectangle. PqsR box: PqsR binding region, RBS: *pqsA* ribosome binding site, TSS1: transcriptional start site 1.

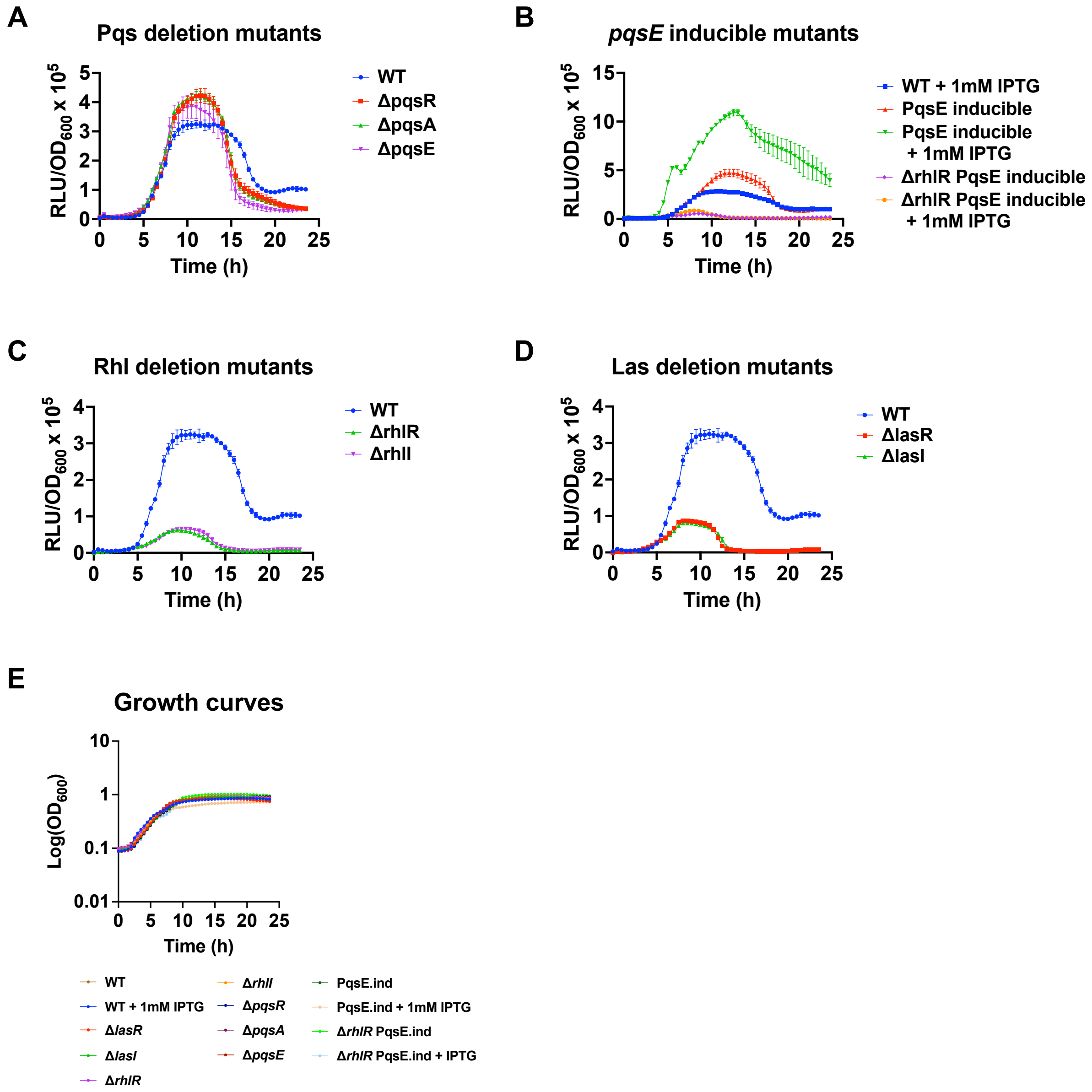


Supplementary Figure 3 Transcriptional activity of the *lrs1* promoter in QS system mutants in LB. A) *lrs1* promoter activity in the WT and the Pqs deletion mutants, B) *lrs1* promoter activity in the WT and the *pqsE* inducible (PqsE.ind) mutants, C) *lrs1* promoter activity in the WT and the Rhl deletion mutants, D) *lrs1* promoter activity in the WT and the Las deletion mutants, E) Growth curves of all strains. Measurements of bioluminescence (relative luminesce units, RLU) were taken every 30 min for 24 h, and were normalised against growth ($OD_{600}$). Error bars represent standard deviation of three biological replicates with three technical replicates.

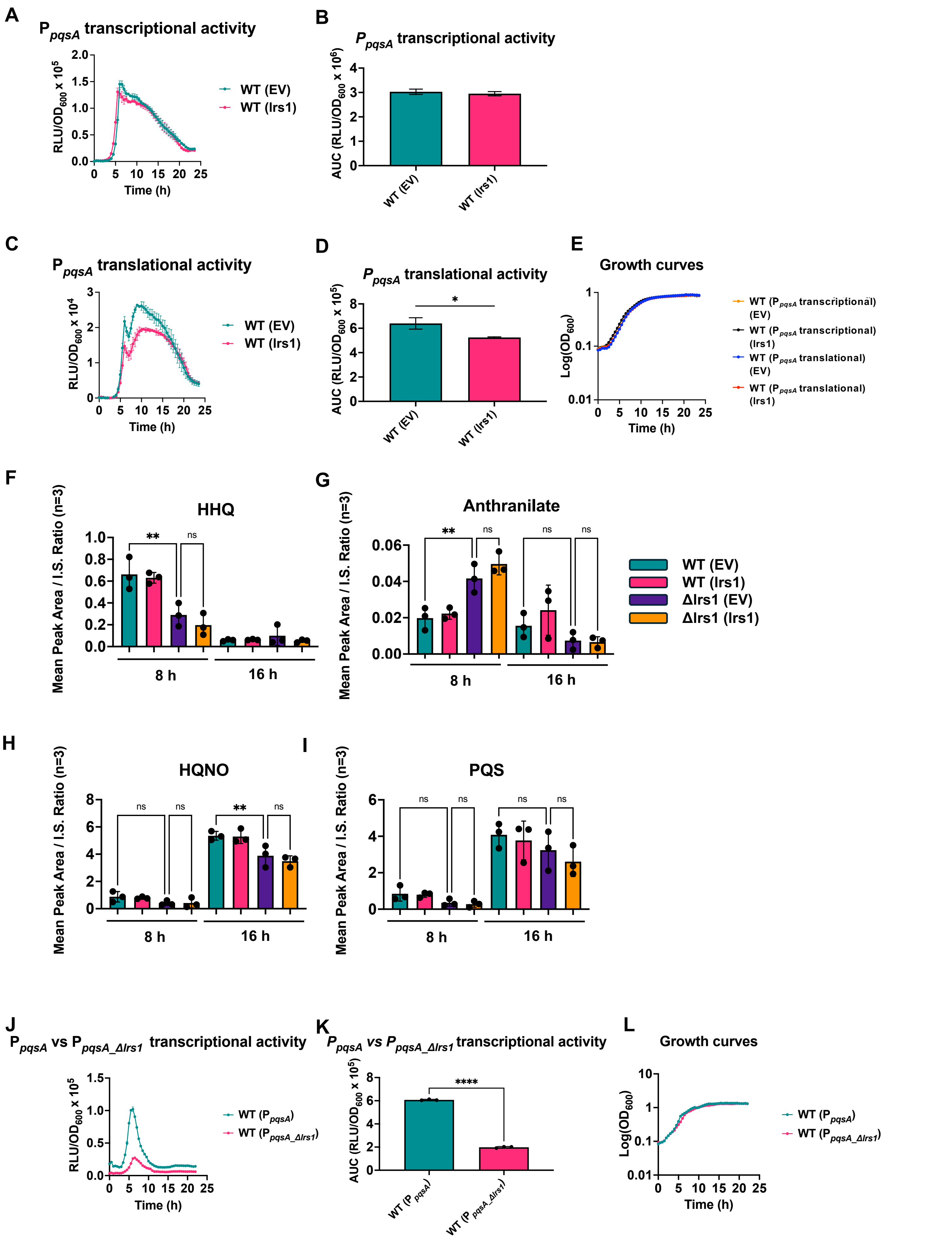

A
PpqsA transcriptional activity
RLU/OD600 x 10^5
Time (h)
WT (EV)
WT (lrs1)
B
PpqsA transcriptional activity
AUC (RLU/OD600 x 10^6)
WT (EV)
WT (lrs1)
C
PpqsA translational activity
RLU/OD600 x 10^4
Time (h)
WT (EV)
WT (lrs1)
D
PpqsA translational activity
AUC (RLU/OD600 x 10^5)
*
WT (EV)
WT (lrs1)
E
Growth curves
Log(OD600)
Time (h)
WT (PpqsA transcriptional) (EV)
WT (PpqsA transcriptional) (lrs1)
WT (PpqsA translational) (EV)
WT (PpqsA translational) (lrs1)
F
HHQ
Mean Peak Area / I.S. Ratio (n=3)
**
ns
8 h
16 h
G
Anthranilate
Mean Peak Area / I.S. Ratio (n=3)
**
ns
8 h
16 h
WT (EV)
WT (lrs1)
Δlrs1 (EV)
Δlrs1 (lrs1)
H
HQNO
Mean Peak Area / I.S. Ratio (n=3)
ns
**
8 h
16 h
I
PQS
Mean Peak Area / I.S. Ratio (n=3)
ns
8 h
16 h
J
PpqsA vs PpqsA_Δlrs1 transcriptional activity
RLU/OD600 x 10^5
Time (h)
WT (PpqsA)
WT (PpqsA_Δlrs1)
K
PpqsA vs PpqsA_Δlrs1 transcriptional activity
AUC (RLU/OD600 x 10^5)
****
WT (PpqsA)
WT (PpqsA_Δlrs1)
L
Growth curves
Log(OD600)
Time (h)
WT (PpqsA)
WT (PpqsA_Δlrs1)

Supplementary Figure 4 Lrs1 overexpression does not impact the $P_{pqsA}$ activity. A) Transcriptional activity of the *pqsA* promoter ($P_{pqsA}$) in the PAO1-L WT with the empty vector (EV) or the *lrs1* overexpressing plasmid (lrs1) in presence of 0.2% L-arabinose. B) Area Under the Curve of the line graph in A. C) Translational activity of the *pqsA* promoter ($P_{pqsA}$) in the PAO1-L WT with the empty vector (EV) or the *lrs1* overexpressing plasmid (lrs1) in presence of 0.2% L-arabinose. D) Area Under the Curve of the line graph in C. E) Growth curves the strains with either transcriptional or translational $P_{pqsA}$ bioluminescent reporter and EV or *lrs1* vectors. Error bars represent standard deviation of three biological replicates with three technical replicates. F) HHQ G) Anthranilate, H) HQNO, I) PQS levels in culture supernatant after 8 h and 16 h of growth in LB + 0.2% L-arabinose. Error bars represent standard deviation of three biological replicates. One-way ANOVA was used for statistical analysis. * : p-value ≤ 0.05, ** : p-value ≤ 0.01. J) Transcriptional activity of the *pqsA* promoter without ($P_{pqsA}$) or with the 138 bp internal deletion of *lrs1* ($P_{pqsA\_\Delta lrs1}$). The two bioluminescent reporters were monitored in WT PAO1-L strains in LB for 24 h. K) Area Under the Curve of the line graph in A. L) Growth curves the strains with the $P_{pqsA}$ or $P_{pqsA\_\Delta lrs1}$ bioluminescent reporters.

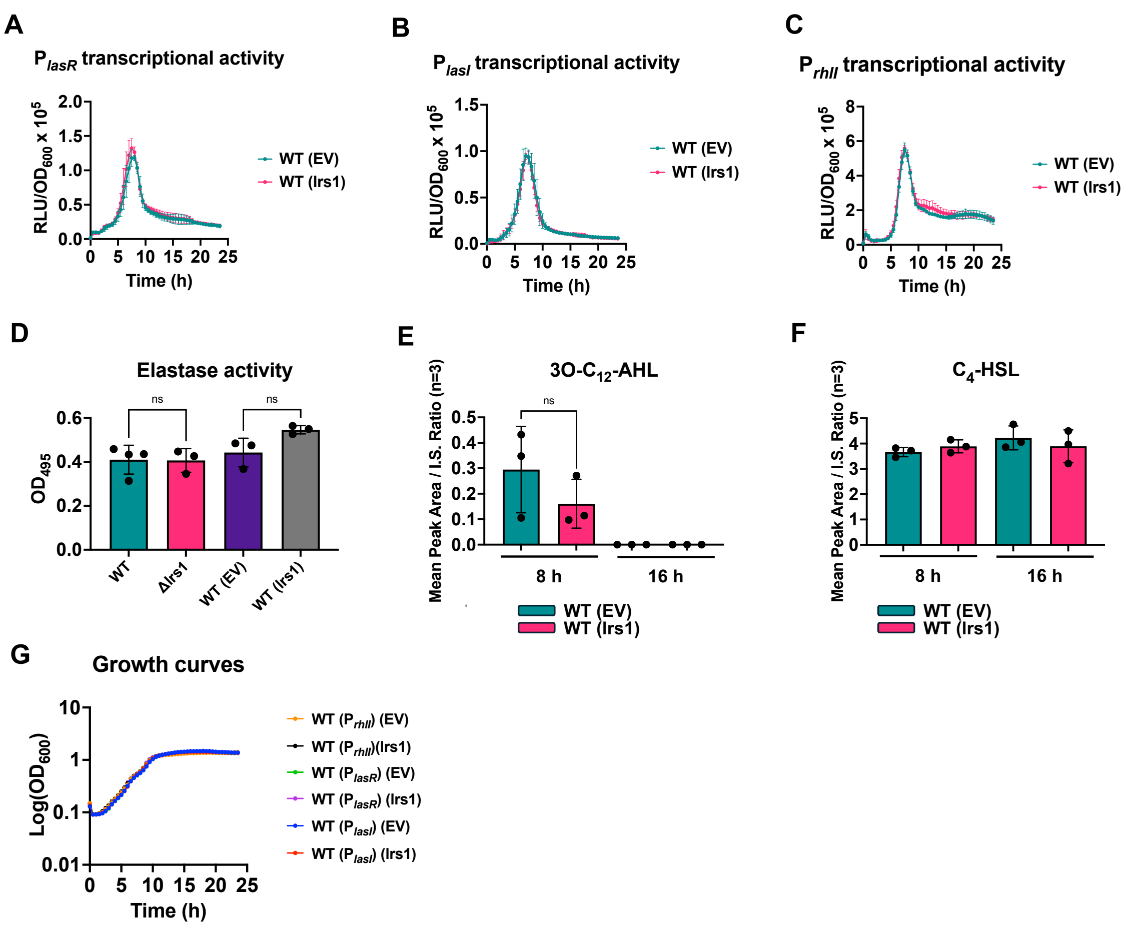


Supplementary Figure 5 Overexpression of Lrs1 does not impact the Las or the Rhl QS systems. A) Transcriptional activity of the *lasR* promoter ($P_{lasR}$) throughout growth in presence of overexpressing *lrs1* or the empty vector, B) Transcriptional activity of the *lasI* promoter ($P_{lasI}$) throughout growth in presence of overexpressing *lrs1* or the empty vector, C) Transcriptional activity of the *rhlI* promoter ($P_{rhlI}$) throughout growth in presence of overexpressing *lrs1* or the empty vector. Measurements of bioluminescence (relative luminesce units, RLU) were taken every 30 min for 24 h, and normalised against growth ($OD_{600}$). Error bars represent standard deviation of three biological replicates with three technical replicates. D) Elastase activity assay in culture supernatants in LB. Samples were taken after 16 h of incubation. Quantification of E) 3O-$C_{12}$-AHL and F) $C_4$-HSL in the WT with the empty vector or the overexpressing *lrs1*, after 8 h and 16 h growth in LB. Error bars represent standard deviation of three biological replicates. G) Growth curves of strains with the $P_{lasR}$, $P_{lasI}$ , $P_{rhlR}$ bioreporter

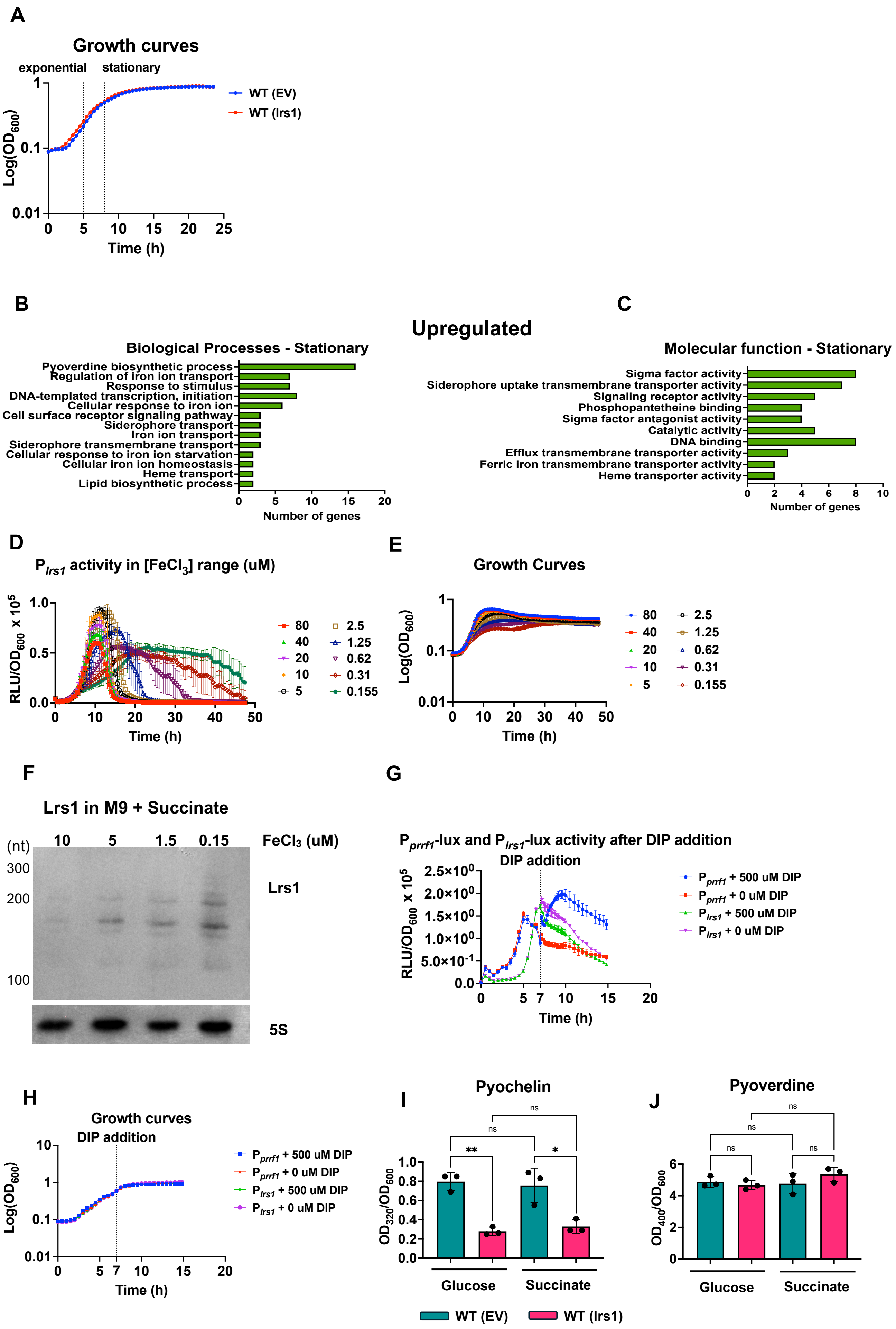
A
Growth curves
exponential
stationary
Log(OD600)
Time (h)
WT (EV)
WT (lrs1)
B
Upregulated
Biological Processes - Stationary
Pyoverdine biosynthetic process
Regulation of iron ion transport
Response to stimulus
DNA-templated transcription, initiation
Cellular response to iron ion
Cell surface receptor signaling pathway
Siderophore transport
Iron ion transport
Siderophore transmembrane transport
Cellular response to iron ion starvation
Cellular iron ion homeostasis
Heme transport
Lipid biosynthetic process
Number of genes
C
Molecular function - Stationary
Sigma factor activity
Siderophore uptake transmembrane transporter activity
Signaling receptor activity
Phosphopantetheine binding
Sigma factor antagonist activity
Catalytic activity
DNA binding
Efflux transmembrane transporter activity
Ferric iron transmembrane transporter activity
Heme transporter activity
Number of genes
D
Plrs1 activity in [FeCl3] range (uM)
RLU/OD600 x 10^5
Time (h)
80 40 20 10 5 2.5 1.25 0.62 0.31 0.155
E
Growth Curves
Log(OD600)
Time (h)
80 40 20 10 5 2.5 1.25 0.62 0.31 0.155
F
Lrs1 in M9 + Succinate
(nt)
10 5 1.5 0.15
FeCl3 (uM)
300
200
100
Lrs1
5S
G
Pprrf1-lux and Plrs1-lux activity after DIP addition
DIP addition
RLU/OD600 x 10^5
Time (h)
Pprrf1 + 500 uM DIP
Pprrf1 + 0 uM DIP
Plrs1 + 500 uM DIP
Plrs1 + 0 uM DIP
H
Growth curves
DIP addition
Log(OD600)
Time (h)
Pprrf1 + 500 uM DIP
Pprrf1 + 0 uM DIP
Plrs1 + 500 uM DIP
Plrs1 + 0 uM DIP
I
Pyochelin
OD320/OD600
ns
**
*
Glucose
Succinate
J
Pyoverdine
OD400/OD600
ns
Glucose
Succinate
WT (EV)
WT (lrs1)

Supplementary Figure 6 Lrs1 is involved in iron uptake. A) Sampling points for RNA-seq of WT with the empty vector (EV) or the *lrs1* overexpressing plasmid (lrs1). Gene ontology enrichment analysis of the upregulated genes in stationary phase for B) Biological processes, C) Molecular function. D) Lrs1 promoter activity in concentration range of $FeCl_3$, from 80 uM down to 0.155 uM. E) Growth curves of WT with the $P_{lrs1}$ bioluminescent reporter in decreasing iron concentrations. Measurements were taken every 30 min for 48 h. Error bars represent standard deviation of three biological replicates with two technical replicates. F) Northern blot of Lrs1 in decreasing concentrations of $FeCl_3$ in M9 with succinate as sole carbon source. 5S rRNA was used as control. G) Transcriptional activity of the bioluminescence reporters $P_{lrs1}$ and $P_{rrf1}$ in PAO1-L WT in LB, after the addition of the iron chelator DIP (bipyridyl) at 7 h post inoculation. H) Growth curves of PAO1-L with the bioluminescence reporters of $P_{lrs1}$ and $P_{rrf1}$ in LB with DIP addition. Quantification of I) Pyochelin and J) Pyoverdine in supernatants of WT with the empty vector and the Lrs1 overexpressing vector. Cells were cultured in M9 with 0.15 uM $FeCl_3$ , 0.2 % L-arabinose and either 2 mM succinate or glucose as carbon source for 20 at which point samples were collected. Error bars represent standard deviation of three biological replicates. Two-tail ANOVA was the statistical test used here. * : p-value ≤ 0.05, ** : p-value ≤ 0.01, ns: no significant difference.

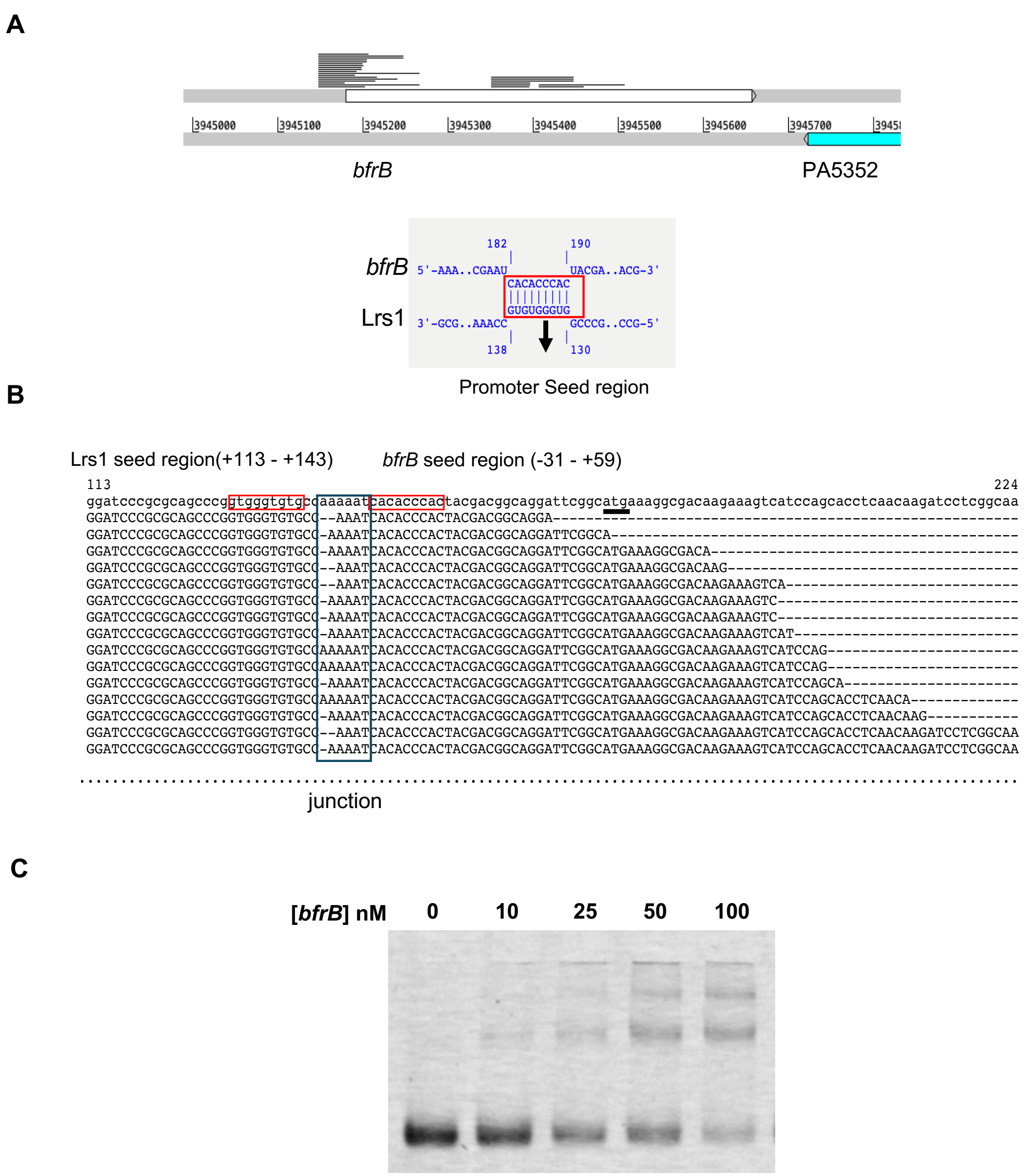


Supplementary Figure 7 Lrs1 interacts with *bfrB in vivo* and *in vitro*. A) Comparison of GRIL-seq targets and RNA-seq differentially regulated genes (DEGs) indicated *bfrB* as a candidate target of Lrs1 related to iron regulation, B) Visualisation of Lrs1-*bfrB* chimeras mapping on the *bfrB* gene (top) and sRNA-mRNA interaction prediction with IntaRNA (bottom). The input region in IntaRNA was -209 nt to +100 nt of *bfrB*. C) Alignment with MAFFT of Lrs1- *bfrB* chimeras. Red rectangles indicate the seed regions and blue rectangular indicate the junction area with putative poly-adelylation. D) EMSA of Lrs1 tagged with the Atto fluorophore and *bfrB* mRNA (-133 nt - +70 nt, +1: *bfrB* translational start site).

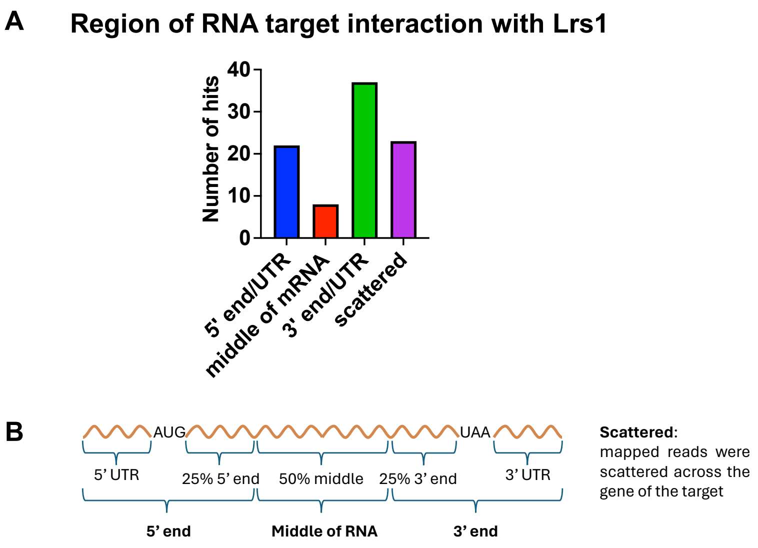


Supplementary Figure 8 Lrs1 interacts with the 5' and the 3' UTR of its target RNAs. A) Column graph representing the positions of the chimeric hits on the RNA bound to Lrs1, B) schematic diagram demonstrating how the different regions of the mRNA target were defined.

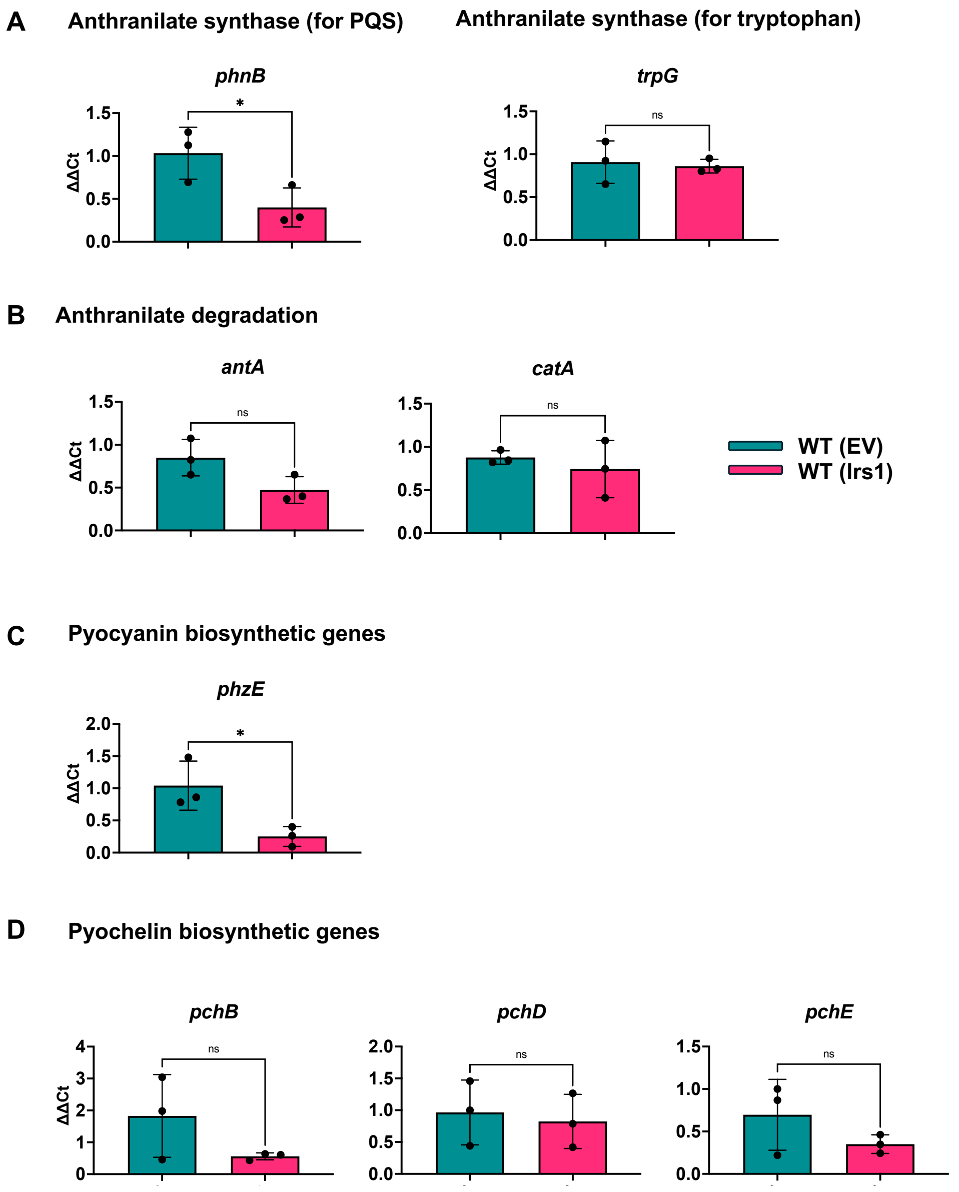


Supplementary Figure 9 RT-qPCRs RT-qPCR analysis of genes participating in A) anthranilate synthesis, B) Anthranilate degradation, C) pyocyanin biosynthesis, D) pyochelin production in the WT and the *lrs1* overexpressing strain at 0.15 μM $FeCl_3$. Cells were grown in M9 minimal media with 0.15 μM $FeCl_3$, 0.2% L-arabinose, and 2 mM glucose for 20 h at which point samples were collected. Error bars represent standard deviation of three biological replicates. T-test was used for statistical analysis. ns: not significant. * : p-value ≤ 0.05, ** : p-value ≤ 0.01.

| **Locus_tag** | **Gene** | **Upregulated in low iron in Ochsner *et al* 2002** | **$\log_2$ Fold Change** | **P-adj** | **ECFσ regulator** |
|---|---|---|---|---|---|
| **Upregulated in stationary phase** | | | | | |
| PA0470 | *fiuA* | - | 1.17 | 5.81E-04 | FiuI |
| PA0472 | *fiuI* | - | 1.15 | 4.76E-03 | FiuI |
| PA0672 | *hemO* | Yes | 1.81 | 8.90E-04 | - |
| PA0707 | *toxR* | Yes | 1.51 | 1.89E-09 | PvdS |
| PA0931 | *pirA* | - | 1.15 | 3.87E-03 | - |
| PA1300 | *hxuI* | Yes | 2.09 | 1.13E-04 | HxuI |
| PA1301 | *hxuR* | Yes | 2.01 | 2.41E-05 | HxuI |
| PA1364 | NA | - | 1.52 | 1.08E-03 | - |
| PA1911 | *femR* | - | 1.62 | 9.50E-04 | FemI |
| PA1912 | *femI* | - | 1.33 | 8.45E-03 | FemI |
| PA2033 | NA | Yes | 2.02 | 2.41E-05 | - |
| PA2034 | NA | Yes | 2.06 | 1.77E-04 | - |
| PA2384 | NA | Yes | 1.70 | 9.83E-04 | - |
| PA2385 | *pvdQ* | Yes | 1.68 | 5.21E-03 | PvdS |
| PA2386 | *pvdA* | Yes | 1.98 | 1.29E-03 | PvdS |
| PA2389 | *pvdR* | Yes | 1.21 | 5.29E-08 | PvdS |
| PA2390 | *pvdT* | - | 1.38 | 1.89E-05 | PvdS |
| PA2391 | *opmQ* | - | 1.09 | 8.73E-04 | PvdS |
| PA2392 | *pvdP* | Yes | 1.91 | 5.34E-07 | PvdS |
| PA2393 | NA | Yes | 1.95 | 4.40E-03 | PvdS |
| PA2394 | *pvdN* | Yes | 2.04 | 4.63E-04 | PvdS |
| PA2396 | *pvdF* | Yes | 1.41 | 9.83E-04 | PvdS |
| PA2397 | *pvdE* | Yes | 2.00 | 3.03E-03 | PvdS |
| PA2399 | *pvdD* | Yes | 1.61 | 2.36E-05 | PvdS |
| PA2400 | *pvdJ* | Yes | 1.74 | 1.37E-05 | PvdS |
| PA2402 | *pvdI* | Yes | 1.70 | 8.16E-05 | PvdS |
| PA2412 | NA | Yes | 1.89 | 3.44E-03 | PvdS |
| PA2413 | *pvdH* | Yes | 1.50 | 1.64E-02 | PvdS |
| PA2424 | *pvdL* | Yes | 1.94 | 2.41E-05 | PvdS |
| PA2425 | *pvdG* | Yes | 1.88 | 5.22E-04 | PvdS |
| PA2426 | *pvdS* | Yes | 2.17 | 5.80E-05 | PvdS |

| | | | | | |
|---|---|---|---|---|---|
| PA2427 | NA | Yes | 1.62 | 2.04E-07 | - |
| PA2451 | NA | Yes | 1.11 | 5.60E-04 | - |
| PA2467 | *foxR* | Yes | 1.20 | 4.76E-03 | FoxI |
| PA2468 | *foxI* | Yes | 1.19 | 3.15E-03 | FoxI |
| PA2531 | NA | Yes | 1.74 | 1.13E-04 | PvdS |
| PA2686 | *pfeR* | - | 1.07 | 2.40E-03 | - |
| PA3407 | *hasAp* | Yes | 1.42 | 1.30E-04 | HasI |
| PA3408 | *hasR* | Yes | 1.44 | 9.83E-04 | HasI |
| PA3410 | *hasI* | Yes | 2.35 | 3.55E-05 | HasI |
| PA3899 | *fecI* | Yes | 1.30 | 3.13E-03 | FecI |
| PA3901 | *fecA* | Yes | 1.65 | 1.52E-03 | FecI |
| PA4158 | *fepC* | Yes | 1.58 | 5.13E-03 | - |
| PA4168 | *fpvB* | - | 1.60 | 5.43E-04 | - |
| PA4227 | *pchR* | Yes | 1.12 | 1.70E-02 | - |
| PA4467 | NA | Yes | 1.72 | 9.02E-04 | - |
| PA4468 | *sodM* | Yes | 2.16 | 4.74E-06 | - |
| PA4469 | N/A | Yes | 2.23 | 1.08E-04 | - |
| PA4470 | *fumC1* | Yes | 2.48 | 2.28E-07 | - |
| PA4471 | NA | | 2.48 | 3.55E-05 | - |
| PA4515 | NA | Yes | 1.09 | 1.51E-02 | - |
| PA4570 | NA | Yes | 2.21 | 2.41E-05 | - |
| PA4704.1 | *prrF1* | - | 2.22 | 7.17E-03 | PvdS |
| PA4708 | *phuT* | Yes | 1.28 | 1.31E-03 | - |
| PA4709 | *phuS* | Yes | 1.73 | 2.25E-03 | - |
| PA4710 | *phuR* | Yes | 1.88 | 8.46E-05 | - |
| PA4896 | NA | Yes | 1.21 | 2.62E-02 | PA4896 |
| PA4897 | NA | Yes | 1.35 | 1.60E-04 | PA4896 |
| **Downregulated in stationary phase** | | | | | |
| PA4222 | NA | Yes | -2.17 | 3.01E-03 | - |
| PA0713 | NA | - | -1.03 | 2.58E-02 | - |
| PA1546 | *hemN* | - | -1.04 | 1.74E-02 | - |
| PA1591 | NA | - | -1.51 | 4.69E-03 | - |
| PA1774 | *crfX* | - | -1.00 | 3.39E-03 | - |
| PA3531 | *bfrB* | - | -1.05 | 1.56E-08 | - |
| PA3720 | NA | - | -1.22 | 5.56E-05 | - |
| PA3721 | *nalC* | - | -1.73 | 4.90E-09 | - |

| | | | | | |
|---|---|---|---|---|---|
| PA4218 | *ampP* | Yes | -2.05 | 3.00E-03 | - |
| PA4219 | *ampO* | - | -2.16 | 1.34E-02 | - |
| PA4221 | *fptA* | Yes | -1.91 | 5.04E-03 | - |
| PA4223 | NA | Yes | -1.67 | 7.71E-03 | - |
| PA4224 | *pchG* | Yes | -2.06 | 2.78E-02 | - |
| PA4225 | *pchF* | Yes | -1.95 | 2.03E-02 | - |
| PA4230 | *pchB* | - | -2.08 | 3.95E-02 | - |
| PA4231 | *pchA* | - | -2.11 | 3.20E-04 | - |
| PA4965 | NA | - | -1.27 | 5.59E-03 | - |
| **Downregulated in exponential phase** | | | | | |
| PA4218 | *ampP* | Yes | -3.78 | 1.95E-09 | - |
| PA4219 | *ampO* | - | -3.84 | 5.74E-07 | - |
| PA4221 | fptA | Yes | -3.68 | 1.18E-10 | - |
| PA4222 | NA | Yes | -2.61 | 6.64E-04 | - |
| PA4223 | NA | Yes | -2.99 | 3.69E-08 | - |
| PA4224 | *pchG* | Yes | -3.13 | 3.39E-04 | - |
| PA4225 | *pchF* | Yes | -3.39 | 5.88E-07 | - |
| PA4226 | *pchE* | - | -3.71 | 2.10E-07 | - |
| PA4228 | *pchD* | Yes | -3.64 | 2.10E-07 | - |
| PA4229 | *pchC* | - | -3.26 | 3.39E-04 | - |
| PA4230 | *pchB* | - | -3.11 | 1.25E-03 | - |
| PA4231 | *pchA* | - | -2.81 | 6.81E-07 | - |
| PA4328 | NA | - | -1.03 | 4.97E-02 | - |

Supplementary Table 1 Differentially regulated genes by Lrs1 overexpression in exponential and stationary growth phases.

| **Locus_tag** | **Gene** | **Description** | **Regions of reads** | **% of total reads** |
|---|---|---|---|---|
| PA3724 | *lasB* | elastase LasB | mainly 3' end, 5' UTR | 14.00 |
| PA5492.1 | *ersA* | sRNA | 3' end half of sRNA | 11.26 |
| PA1431 | *rsaL* | regulatory protein RsaL | equally separated at 3' and 5' end of gene | 4.26 |
| PA0958 | *oprD* | Basic amino acid, basic peptide and imipenem outer membrane porin OprD precursor | 5' end and middle of the gene | 4.57 |
| PA0852 | *cbpD* | chitin-binding protein CbpD precursor | 3' end | 3.96 |
| PA0865 | *hpd* | 4-hydroxyphenylpyruvate dioxygenase | mainly 3' end, 5' end | 10.65 |
| PA1032 | *quiP* | Acyl-homoserine lactone (AHL) acylase PvdQ | 3' end | 2.28 |
| PA1802 | *clpX* | ATP-dependent protease Clp, ATPase subunit | 3' end | 10.50 |
| PA3305.1 | *phrS* | sRNA | 5' end half of sRNA | 7.76 |
| PA0447 | *gcdH* | glutaryl-CoA dehydrogenase | scattered | 1.67 |
| PA1831 | - | Broad specificity phosphatase PhoE | middle of the gene | 2.44 |
| PA2069 | - | probable carbamoyl transferase | scattered | 1.37 |
| PA2250 | *lpdV* | lipoamide dehydrogenase-Val | scattered | 2.59 |
| PA2371 | - | ATP-dependent Clp protease ATP-binding subunit ClpA (ClpV3) | 3' end | 1.22 |
| PA2620 | *clpA* | ATP-binding protease component ClpA | 3' end | 1.37 |
| PA3105 | *xcpQ* | general secretion pathway protein D | 3' end | 1.22 |
| PA3162 | *rpsA* | 30S ribosomal protein S1 | scattered | 3.04 |
| PA3479 | *rhlA* | rhamnosyltransferase chain A | scattered | 2.44 |
| PA4420 | - | HP | scattered | 1.98 |
| PA4607 | - | HP | scattered | 2.89 |
| PA4922 | *azu* | azurin precursor | 3' end | 2.44 |
| PA5170 | *arcD* | arginine/ornithine antiporter | 3' end | 2.28 |
| PA1150 | *pys2* | pyocin S2 | 3' end | 3.81 |

Supplementary Table 2 GRIL-seq targets of Lrs1 identified in both exponential and stationary conditions

| Locus_tag | Gene name | Description | Regions of reads | % of total reads |
|---|---|---|---|---|
| PA4217 | *phzS* | flavin-containing monooxygenase | mainly 3' end, 5' UTR | 5.73 |
| PA4315 | *mvaT* | transcriptional regulator MvaT | mainly 5' UTR, 3' end | 4.78 |
| PA2016 | *liuR* | regulator of *liu* genes | 3' end half of the gene | 4.54 |
| PA3190 | - | probable binding protein component of ABC sugar transporter | 3' end | 4.06 |
| PA5307 | - | HP | 3' end | 3.46 |
| PA3187 | - | probable ATP-binding component of ABC transporter | 5' end, 3' UTR | 3.11 |
| PA4463 | - | HP | 5' UTR, 3' end | 2.87 |
| PA4277 | *tufB* | elongation factor Tu | scattered | 2.15 |
| PA4584 | - | HP | middle of the gene | 2.63 |
| PA4741 | *rpsO* | 30S ribosomal protein S15 | 3' end | 1.91 |
| PA0995 | *ogt* | methylated-DNA--protein-cysteine methyltransferase | middle of the gene | 2.15 |
| PA1777 | *oprF* | Major porin and structural outer membrane porin OprF precursor | mainly 5' UTR, 5' end | 3.58 |
| PA3152 | *hisH2* | glutamine amidotransferase | middle of the gene | 1.67 |
| PA3531 | *bfrB* | bacterioferritin | 5' UTR | 2.87 |
| PA3533 | - | Glutaredoxin-related protein GrxD | 3' end | 2.15 |
| PA4272 | *rplJ* | 50S ribosomal protein L10 | 5' end | 1.67 |
| PA3186 | *oprB* | Glucose/carbohydrate outer membrane porin OprB precursor | 3' end, 5' UTR | 3.94 |
| PA2743 | *infC* | translation initiation factor IF-3 | scattered | 1.31 |
| PA3385 | *amrZ* | alginate and motility regulator Z | mainly 3' end, 5' end | 1.91 |
| PA4131 | - | probable iron-sulfur protein | middle of the gene | 1.79 |
| PA4266 | *fusA1* | elongation factor G | 3' end | 1.43 |
| PA4726.11 | *crcZ* | sRNA | 5' end half of gene | 2.39 |
| PA5171 | *arcA* | arginine deiminase | scattered | 1.43 |
| PA5240 | *trxA* | thioredoxin | scattered | 1.43 |

| | | | | |
|---|---|---|---|---|
| PA5557 | *atpH* | ATP synthase delta chain | 5' end | 1.55 |
| PA0762 | *algU* | sigma factor AlgU | 3' end | 1.31 |
| PA2291 | - | probable glucose-sensitive porin | 3' end | 1.55 |
| PA0122 | - | probable hemolysin RahU | middle of the gene | 1.79 |
| PA0826.2 | *ssrA* | tmRNA | split in the 3' end and 5' end of the gene | 1.67 |
| PA1372 | - | HP | 3' end | 1.19 |
| PA1584 | *sdhB* | succinate dehydrogenase (B subunit) | 5' end | 1.19 |
| prc (PA3257) | *prc* | Periplasmic tail-specific protease AlgO | 3' end | 1.43 |
| PA4554 | *pilY1* | type 4 fimbrial biogenesis protein PilY1 | scattered | 1.43 |
| PA4740 | *pnp* | polyribonucleotide nucleotidyltransferase | scattered | 1.43 |
| PA4751 | *ftsH* | cell division protein FtsH | scattered | 1.19 |
| PA0588 | - | HP | scattered | 1.67 |
| PA2445 | *gcvP2* | glycine cleavage system protein P2 | scattered | 1.91 |
| PA0977 | - | HP | 3' UTR of tRNA-lys | 1.19 |
| PA1901 | *phzC2* | phenazine biosynthesis protein PhzC | scattered | 1.43 |
| PA2224 | - | HP | 3' end | 1.19 |
| PA4265 | *tufA* | elongation factor Tu | scattered | 1.19 |
| PA4726.2 | - | P30 sRNA transcribed from complementary strand of *crcZ* | on CrcZ* | 1.31 |
| PA0084 | - | TssC1 | scattered | 1.19 |
| PA0038 | - | HP | 3' end | 1.19 |
| PA1000 | *pqsE* | Quinolone signal response protein | scattered | 1.19 |
| PA1800 | *tig* | FKBP-type peptidyl-prolyl cis-trans isomerase (trigger factor) | 3' end | 1.19 |
| PA4212 | *phzC1* | phenazine biosynthesis protein PhzC | scattered | 1.43 |
| PA4385 | *groEL* | GroEL protein | scattered | 1.19 |
| PA4761 | *dnaK* | DnaK protein | 5' end, middle of the gene | 1.91 |

6 Supplementary Table 3 GRIL-seq targets of Lrs1 identified in exponential samples only.

| Locus_tag | Gene | Description | Regions of reads | % of total reads |
|---|---|---|---|---|
| PA1874 | - | HP | scattered | 48.94 |
| PA1041 | - | probable outer membrane protein precursor | 3' end | 10.64 |
| PA2030 | - | HP | 3' end | 30.85 |
| PA4785 | - | probable acyl-CoA thiolase | 3' end | 9.57 |

Supplementary Table 4 GRIL-seq targets of Lrs1 identified in stationary samples only.

| Strains | Description | Reference |
|---|---|---|
| DH5α | $F^-$*φ80ΔlacZM15 Δ(lacZYA-argF)U169 recA1 hsdR17 (r-km+k) supE44 thi-1 relA1 gyrA96* | (Grant *et al.*, 1990) |
| S17-1 λ*pir* | *thi pro hsdR*$^-$ *M*$^+$ *recA* RP4-2-Tc::Mu-Km::Tn7 λ*pir* *Tp*$^R$ *Sm*$^{R-}$ | (de Lorenzo *et al.*, 1993) |
| PAO1-L | Lausanne subline of wild type *P. aeruginosa* PAO1 strain | (Dubern *et al.*, 2022) |
| PAO1-L Δ*lasR* | PAO1-L mutant with *lasR* in frame deletion | This study |
| PAO1-L Δ*lasI* | PAO1-L mutant with *lasI* in frame deletion | This study |
| PAO1-L Δ*rhlR* | PAO1-L mutant with *rhlR* in frame deletion | This study |
| PAO1-L Δ*rhlI* | PAO1-L mutant with *rhlI* in frame deletion | This study |
| PAO1-L Δ*pqsR* | PAO1-L mutant with *pqsR* in frame deletion | This study |
| PAO1-L Δ*rhlR* pqsE.ind | PAO1-L with *pqsE* under the $P_{tac}$ inducible promoter and rhlR in frame deletion | This study |
| PAO1-L Δ*pqsA* | PAO1-L mutant with *pqsA* in frame deletion | This study |
| PAO1-L PqsE.ind | PAO1-L with *pqsE* under the $P_{tac}$ inducible promoter | (Rampioni *et al.*, 2010) |
| PAO1-L Δlrs1 | PAO1-L mutant with 138 nt *lrs1* deletion from -337 bp to -200 bp relative to *pqsA* start codon | This study |
| PAO1-L Δ*pqsE* | PAO1-L mutant with *pqsA* in frame deletion | This study |

Supplementary Table 5 Strains used in this study.

| Name | Description | Reference |
|---|---|---|
| pminiCTXlux Tc | Site-specific integration vector for *P. aeruginosa*; $Tc^R$ | (Hoang *et al.*, 2000) |
| pminiCTXlux | Site-specific integration vector for *P. aeruginosa*; the gentamicin resistance gene has been cloned in the *tetR* gene in the unique EagI site to make the vector $Gm^R$ | lab collection |
| pBluescript | Cloning vector; ColE1 replicon; $Amp^R$ | Agilent Technologies, US |
| pTS1 | pME3087 (Suicide vector; ColE1 replicon, IncP-1, Mob; $Tc^R$) modified with *sac*B counter-selection and an expanded multiple cloning site, $Tc^R$ | (Scott *et al.*, 2017) |
| pME6032 | pVS1-p15A shuttle expression (IPTG-inducible) vector, $Tc^R$ | (Heeb, Blumer and Haas, 2002) |
| pME3087 | Suicide vector; ColE1 replicon, IncP-1, Mob; $Tc^R$ | (Voisard *et al.*, 1994) |
| pBS_1 | pBS carrying -615 to +357 area of $P_{pqsA}$ (+1 *pqsA* translational start site | This study |
| pBS1_mut | pBS_1 derivative, reversed amplified and self-ligated to make the substitution of the CGTTC upstream of TSS1 with AAGAA. | This study |
| pDP013 | pTS1 suicide vector to make Δ*lrs1* deletion | This study |
| pEX18::Δpqs R | Gene replacement vector oriT+, sacB; for in-frame deletion of *pqsR* in PAO1-L, $Gm^R$ | (Ilangovan *et al.*, 2013) |
| pKH6 | pBBR1 derivative, with $P_{bad}$-AraC L-arabinose inducible system for transcription of sRNAs. $Gm^R$ | (Han *et al.* 2016) |
| pKH13-t4rnl1 | pKH13 vector with IPTG inducible $P_{tac}$ promoter for the induction of T4 RNA ligase gene. $Cm^R$ | (Han *et al.*, 2016) |
| pKH6-lrs1 | pKH6 carrying a 248 bp *lrs1* insertion between XbaI and EcoRI for transcription from the pBAD promoter. | This study |
| $P_{lrs1}$-lux | pminiCTXlux with a 534 *lrs1* promoter transcriptionally fused between HindIII and PstI, | This study |
| $P_{pqsA}$-lux | $P_{pqsA}$ transcriptional fusion in pminiCTXluxTc, at PstI and HindIII sites. $Tc^R$ | This study |
| $P_{lasR}$-luxTc | pminiCTXluxTc with the lasR 523 bp promoter transcriptionally fused between HindIII and PstI. $Tc^R$ | This study |
| $P_{lasI}$-luxTc | pminiCTXluxTc with the 243 bp *lasI* promoter transcriptionally fused between HindIII and PstI. $Tc^R$ | This study |
| $P_{rhlI}$-luxTc | pminiCTXluxTc with the 451 bp *rhlI* promoter transcriptionally fused between HindIII and PstI. $Tc^R$ | This study |
| $P_{pqsA}$::luxTc | pminiCTXluxTc with $P_{pqsA}$ -495 to +5 translational fusion at the XcmI. | This study |
| $P_{pqsA}$_Δlrs1::l uxTc | pminiCTXluxTc with $P_{pqsA}$ (-495 to +5) with a 138bp *lrs1* deletion translational fusion at the XcmI. $Tc^R$ | This study |
| pBx-Spas-sgRNA-Gm | Plasmid containing S. pasteurianus sgRNA with no 20bp target (Empty control) Gentamycin marker for *P. aeruginosa*. $Gm^R$ | Acquired from Addgene. (Tan, Reisch and Prather, 2018) |
| pUC18-miniTn7-Plac-dCas9-Gm- | Tn7 integrating plasmid with S. pasteurianus dCas9 expressed from the Plac promoter for expression in *P. aeruginosa*. $Gm^R$ | Acquired from Addgene. (Tan, Reisch and Prather, 2018) |
| pBx-rne1 | pBx-Spas-sgRNA-Gm with the guide RNA for the region of | This study |
| pME6032-lasR | IPTG-inducible expression of *lasR* from the $P_{tac}$ promoter | This study |
| pME6032-rhlR | IPTG-inducible overexpression of *rhlR* from the $P_{tac}$ promoter | This study |

| | | |
|---|---|---|
| pME3087-rhlR | pME3087 suicide vector to construct ΔrhlR deletion | This study |
| pME3087-rhlI | pME3087 suicide vector to construct ΔrhlI deletion | This study |
| pME3087-lasR | pME3087 suicide vector to construct ΔlasR deletion | This study |
| pME3087-lasI | pME3087 suicide vector to construct ΔlasI deletion | This study |
| pME3087-pqsE | pME3087 suicide vector to construct ΔpqsE deletion | This study |

Supplementary Table 6 Plasmids used in this study.

| Name | Sequence 5' → 3' | Description |
|---|---|---|
| PAPQF | ACCTTGGAAGTCGAGCGGGAGT | Sequence *pqsA* promoter from pBS_1 |
| PAPQR | CAGGCTGTGCATCACCCCCTTG | Sequence *pqsA* promoter |
| LRS1RV | TATCTGCAGCGCCAAAGGACCGCTCAG | Amplify *pqsA* promoter 987 bp |
| LRS1FV | TATGAATTCGCTACCAGGGTCAGTGTCC | Amplify *pqsA* promoter 987 bp |
| PDP002F | GCGTTGTCGGGAAGATGCGTGA | Sequencing insert in pTS1, PCR after conjugation |
| PDP002R | CGGTCTGCTTCTTCCAGCCCTC | Sequencing insert in pTS1, PCR after conjugation |
| Northlrs1FW | GCCATCTCATGGGTTCGGACGAGG | 202bp *lrs1* template for northern blot |
| Northlrs1T7RV | TAATACGACTCACTATAGGGGGGGAGAAGAACCACGGGCT | 202bp *lrs1* template for northern blot, T7 promoter underlined |
| LRS1PROHINDIIIF | TATAAAGCTTCGGGTCTCGCCGAATGGAATGG | Amplify *lrs1* promoter 534 bp |
| LRS1PROMPSTIR | TATCTGCAGTCGTCCGAACCCATGAGATGGCA | Amplify *lrs1* promoter 534 bp |
| PQSAHINDIIIFW | TATAAGCTTGCTACCAGGGTCAGTGTCC | Insertion of $P_{pqsA}$ in pminiCTXlux between HindIII and EcoRI sites |
| PQSAPSTIRV | TATCTGCAGAGGCGGAACAGAACCTCGGTCA | Insertion of $P_{pqsA}$ in pminiCTXlux between HindIII and EcoRI sites |
| PlasRHindIIIFW | AAAAAAGCTTCGCCGAACTGGAAAAGTGGCTA | make transcriptional fusion of the *lasR* promoter 520 bp to pminiCTXlux |
| PlasRPstIRV | AAAACTGCAGACCGTCAACCAAGGCCATAGC | make transcriptional fusion of the *lasR* promoter 520 bp to pminiCTXlux |
| PlasIHindIIFW | AAAAAAGCTTAAGTTTCCTGGCTTTCCCGTCG | make transcriptional fusion of the *lasI* promoter 242 bp to pminiCTXlux |
| PlasIPstIRV | AAAACTGCAGGCGCCGACCAATTTGTACGATC | make transcriptional fusion of the *lasI* promoter 242 bp to pminiCTXlux |
| PPqsA-translational-F | ATAGTTAAACAGCAACTTAAGTTGAAATTACCCCCATTAACTGCAAATGGCAGGCGAG | to make translational fusion of $P_{pqsA}$ to pminiCTXlux |
| PPqsA-translational-R | AATAATGAATGAAATTTTTTTAGTCATATTTGCCATCCATGACATGACAGAACGTTCCCTCT | to make translational fusion of $P_{pqsA}$ to pminiCTXlux |
| LRS1XBAIFW | AAAATCTAGAGAGCCATCTCATGGGTTCGGACG | make 248 bp *lrs1* complementation in pKH6 |
| LRS1_ECORI_M_RV | AAAAGAATTCCGCTGGGCGAAGCGCGA | make 248 bp *lrs1* complementation in pkh6 |
| Lrs1 P1 | AACCGTTACAACCCTTGCTC | RT of *lrs1* GRIL-Seq samples |
| Lrs1 P2 | CCTCGTCCGAACCCATGAG | PCR of *lrs1* RT with P1 primer |
| Lrs1 P3 | TTTGGATCGCGCCGATTGC | PCR of *lrs1* RT with P1 primer |
| Lrs1 poly-A | AAAAAAAAAAAAAAAAAAGGCCTCGTCCGAACCCATGAGAT | Pull-out *lrs1* chimeric RNAs from the GRIL-Seq *lrs1* samples |
| Lrs1 qRT FW | CTTGCTTGGTTGCCGTTCTC | qPCR of Lrs1 79 bp product |
| Lrs1 qRT RV | CAATCGGCGCGATCCAAAC | qPCR of Lrs1 79 bp product |
| rpoD-F | GGGCGAAGAAGGAAATGGTC | qPCR of endogenous control *rpoD* 178 bp product |
| rpoD-R | CAGGTGGCGTAGGTGGAGAA | qPCR of endogenous control *rpoD* 178 bp product |
| F_q_t4rnl1+65 | CTCAGATGATGTAAGTGCATCTGGAAG | qPCR of *t4rnl1 mRNA*, product: 109 bp. From (Han *et al.* , 2016) |
| F_q_t4rnl1+171 | CATAATTCCACGACATTCTAGTGCATC | qPCR of*t4rnl1 mRNA*, product: 109 bp. From (Han *et al.*, 2016) |
| lrs1190delF1FW | TCTAGAGTCGACCTGCACCAAAGGACCGCTCAGC | To make *lrs1* deletion 138 bp, fragment 1 500 bp |
| lrs1190delF2RV | AAGCTTGCATGCCTGCATCCGTATAGCTGCGGGT | To make *lrs1* deletion 138 bp, fragment 2 525 bp |
| lrs1138delF2FW | AACCGCGAGAAATTTGGCAAACCAATGACAACCC | To make *lrs1* deletion 138 bp, fragment 2 |
| lrs1138delF1RV | GGGTTGTCATTGGTTTGCCAAATTTCTCGCGGTTTG | To make *lrs1* deletion 148 bp, fragment 1 |
| T7_lrs1_RV | TTTTTTTTCCCCCCCCCGCTGGGCGAAGCGCGATAT | EMSA, amplify 248 bp lrs1 template without ATTO complementary sequence |
| T7_lrs1_RV_with_atto | TTTTTTTTCCCCCCCCCGCTGGGCGAAGCGCGATAT | EMSA, amplify 248 bp Lrs1 template with ATTO complementary sequence, |

| | | |
|---|---|---|
| | | ATTO complementary sequence underlined |
| T7_lrs1_FW | TAATACGACTCACTATAGGGGCCATCTCATGGGTTCGGACGAGG | EMSA, T7 promoter underlined |
| Lrs1_T7_short_atto_RV | TTTTTTTTCCCCCCCCCGGGCTTCGTAGGCCGCG | To make 190 nt version of Lrs1, ATTO complementary sequence underlined |
| T7_bfrB_RV | GGTTGATCGCGATCAGCTCGTT | EMSA |
| T7_bfrB_FW | TAATACGACTCACTATAGGGCGGCAGAACGCGGAAAGAAAAC | EMSA, T7 promoter underlined |
| atto probe | ATTO700-TTTTTTTTCCCCCCCCC | fluorescent probe for EMSA with the ATTO fluorophore covalently conjugated at the 5' (ordered from Eurofins) |
| sgRNA_1_RV | AAACGCGTCGAACCCAGCCTCG | to make double strand DNA of sgRNA for the *rne* promoter in pBx-Spas-sgRNA-Gm |
| sgRNA_1_FW | GCACTTCGAGGCTGGGTTCGAC | to make double strand DNA of sgRNA for the *rne* promoter in p pBx-Spas-sgRNA-Gm |
| LasR-C-F | TATGAATTCATGGCCTTGGTTGACGG | Cloning of *lasR* in pME6032 |
| LasR-C-R | ATAGGTACCTGGGTCTTATTACTCTCTGA | Cloning of *lasR* in pME6032 |
| RhlR-C-F | TATGAATTCATGAGGAATGACGGAGGCT | Cloning of *rhlR* in pME6032 |
| RhlR-C-R | ATAGGTACCTCAGATGAGACCCAGCG | Cloning of *rhlR* in pME6032 |
| FWpqsRUp | ATAAAGCTTTCTTAGAACCGTTCCTGG | To construct the pEX18::ΔpqsR plasmid |
| RVpqsRUp | ATAGGATCCAGGTTATGAATAGGCATC | To construct the pEX18::ΔpqsR plasmid |
| FWpqsRDown | ATAGGATCCAGAGTAGAGCGTTCTCCA | To construct the pEX18::ΔpqsR plasmid |
| RVpqsRDown | ATATCTAGAAGTTCTGCCTGCTCGGCG | To construct the pEX18::ΔpqsR plasmid |
| RhlI-Up-F1 | ATAGAATTCTTTCCGTGGCGCGCGACCAG | Forward primer for RhlI flanking region with EcoRI at the 5' end. |
| RhlI-Up-R1 | ATATCTAGACAAGTCCCCGTGTCGTGCCG | Reverse primer for RhlI flanking region with XbaI site at the 5' end. |
| RhlI-Dn-F1 | ATATCTAGATACCACCCGGAATGGCTGCA | Forward primer for RhlI Down flanking region with XbaI site at the 5' end. |
| RhlI-Dn-R1 | ATAAAGCTTTGGCGCTCCAGGTTGATCGA | Reverse primer for downstream flanking region of RhlI with a HindIII site at the 5' end. |
| RhlR-Up-F1 | ATAGGTACCCCTCGGCGCGCGTGGGATCT | Forward primer for upstream RhlR flanking region with KpnI site at the 5' end. |
| RhlR-Up-R1 | ATATCTAGATGGCCCGGGGTATGACGCTG | Reverse primer for upstream RhlR flanking region with an XbaI site at the 5' end. |
| RhlR-Dn-F1 | ATATCTAGAATCCACCACAAGAACATCCAG | Forward primer for downstream RhlR flanking region with an XbaI site at the 5' end. |
| RhlR-Dn-R1 | ATAAAGCTTAAGACGTCCTTGAGCAGGTA | Reverse primer for downstream RhlR flanking region with a HindIII site at the 5' end. |
| LasR 1 For | ATAAAGCTTCCCTCTAGGACGGGTATCGT | Forward primer for Upstream LasR deletion construct with a HindIII at the 5' end. |
| LasR 1 Rev | ATAGAATTCTGAAGGCGTTCTCGTAGTCC | Reverse primer for Upstream LasR deletion construct with an EcoRI site at the 5' end. |
| LasR 2 For | ATAGAATTCGGGAGAAGGAAGTGTTGCAG | Forward primer for Downstream LasR deletion construct with an EcoRI site at the 5' end. |
| LasR 2 Rev | ATATCTAGAGAGAGAACACAGCCCCAAAA | Reverse primer for Downstream LasR dlgetion construct with a XbaI site at the 5' end. |
| PqsE-UP-F | ATATCTAGAGCGCCGGCGAGAGTCT | Forward primer for Upstream *pqsE* deletion construct with a XbaI at the 5' end. |
| PqsE-UP-R | ATAGGATCCAGCCGAAAGCCTCAACATGG | Reverse primer for Upstream *pqsE* deletion construct with an BamHI site at the 5' end. |
| PqsE- DS- F | ATAGGATCCGACTGAGACGGCACATCCA | Forward primer for Downstream *pqsE* deletion construct with an BamHI site at the 5' end. |
| PqsE- DS -R | ATAGGTACCAGGCTGGACAGGCCATGC | Reverse primer for Downstream *pqsE* digetion construct with a KpnI site at the 5' end. |

3 Supplementary Table 7 Oligos used in this study.
